# HelD is a Global Transcription Factor Enhancing Gene Expression in Rapidly Growing Mycobacteria

**DOI:** 10.1101/2024.11.27.625628

**Authors:** Viola Vaňková Hausnerová, Dilip Kumar, Mahmoud Shoman, Marek Schwarz, Martin Modrák, Jitka Jirát Matějčková, Silvia Neva, Jarmila Havelková, Michaela Šiková, Debora Pospíšilová, Petr Halada, Hana Šanderová, Jana Holubová, Matúš Dohál, Martin Převorovský, Ondřej Staněk, Zdeněk Knejzlík, Věra Dvořáková, Jarmila Hnilicová

## Abstract

HelD protein, also named HelR (encoded by *MSMEG_2174* in *Mycobacterium smegmatis*), interacts with mycobacterial RNA polymerase (RNAP) and affects rifampicin resistance in *Mycobacterium abscessus*. Here, we provide data on rifampicin resistance and *helD* presence in the genomes of other clinically relevant nontuberculous mycobacteria.

We show that *helD* is primarily found in rapidly growing mycobacteria, such as *M. smegmatis*, where we detected HelD at a subset of promoters that can also associate with CarD and RbpA. Transcriptome analysis of a *helD* deletion strain using RNA-seq revealed that HelD enhances gene expression during exponential growth and decreases it in stationary phase, during which we observed reduced levels of CarD, RbpA, and GTP, the initiation nucleotide for the majority of *M. smegmatis* transcripts. We propose a model in which HelD releases abortive RNAP complexes and confirm that HelD dissociates RNAP from the promoter *in vitro*.

HelD not only helps mycobacteria overcome rifampicin treatment but also supports efficient transcription during rapid growth, which indicates a dual role of this transcription regulator.

## INTRODUCTION

Bacterial transcription is an essential process for bacterial growth and viability. A key enzyme involved in transcription is RNA polymerase (RNAP), which is composed of several subunits (β, β′, ω and two αs) forming the RNAP core (Burgess *et al*, 1969). The RNAP core associates with σ factors that target RNAP to gene promoters and help to initiate transcription (Browning & Busby, 2016). In addition to σ factors, mycobacterial RNAP binds to the general transcription factors CarD and RbpA (Stallings & Glickman, 2011; Stallings *et al*, 2009; Tabib-Salazar *et al*, 2013)). CarD and RbpA associate with the RNAP during transcription initiation and stabilize the open promoter complex (Hu *et al*, 2012; Hubin *et al*, 2017a; Rammohan *et al*, 2016; Rammohan *et al*, 2015; Srivastava *et al*, 2013; Stallings *et al*., 2009). Recently, HelD was identified as a protein that binds to non-transcribing RNAP and can interact with RNAP in complex with RbpA and the primary mycobacterial σ factor, σ^A^ (Kouba *et al*, 2020; Kovaľ *et al*, 2024).

HelD is an RNA-polymerase binding protein that was first discovered in 2011 in *Bacillus subtilis* (gene name *helD*, originally *yvgS,*) (Delumeau *et al*, 2011; Wiedermannová *et al*, 2014) and later also in *Mycobacterium smegmatis* (gene name *MSMEG_2174*) (Kouba *et al*., 2020). Due to its role in rifampicin resistance, HelD was named HelR in *Mycobacterium abscessus* and *Streptomyces venezuelae* (Hurst-Hess *et al*, 2022; Surette *et al*, 2022). For simplicity, we will refer to it as HelD. HelD-like proteins are widely distributed among eubacteria, including both Gram positive *Firmicutes* and *Actinobacteria* and Gram negative *Deltaproteobacteria* and *Bacteroides* (Larsen *et al*, 2021). Some actinobacterial species, such as *Frankia* and *Streptomycetes*, which are industrially important producers of antibiotics and other secondary metabolites, have up to five different *helD* genes encoded in their genomes (Larsen *et al*., 2021).

HelD belongs among the superfamily 1 (SF1) helicases, the largest class of helicases with members that participate in many steps of DNA and RNA metabolism (Raney *et al*, 2013). Although HelD shares some conserved sequence motifs with SF1 helicases and is able to hydrolyze ATP and also GTP (Kouba *et al*., 2020), HelD lacks DNA binding domain (Surette *et al*., 2022) and helicase activity has never been detected for HelD (Wiedermannová *et al*., 2014). Instead, HelD proteins were shown to associate with the bacterial RNAPs in *B. subtilis*, *M. smegmatis*, *M. abscessus* and *S. venezuelae* and to release RNAP from DNA (Hurst-Hess *et al*., 2022; Kouba *et al*., 2020; Surette *et al*., 2022; Wiedermannová *et al*., 2014). Based on *in vitro* data, HelD was proposed to promote RNAP recycling after transcription termination, disassemble stalled transcription elongation complexes and inhibit non-specific binding of RNAP to DNA (Kouba *et al*., 2020; Wiedermannová *et al*., 2014).

HelD structures in complex with RNAP in *B. subtilis* and *M. smegmatis* have been solved. Both HelD proteins have N-terminal domains that insert into the RNAP secondary channel (Kouba *et al*., 2020; Newing *et al*, 2020; Pei *et al*, 2020). The secondary channel of RNAP is necessary for the access of nucleoside triphosphates or transcription factors (such as GreA (Laptenko *et al*, 2003; Rutherford *et al*, 2007) or DksA (Lennon *et al*, 2012; Ross *et al*, 2016)) to the active site of RNAP. However, the N-terminal domain of mycobacterial HelD is shorter and does not reach to the RNAP catalytic site, but rather stabilizes the interaction between HelD and RNAP (Kouba *et al*., 2020). The HelD-specific domain enters the RNAP primary channel, where DNA binds during transcription. Consequently, this widens the RNA exit channel of RNAP, forcing both DNA and RNA to leave RNAP (Kouba *et al*., 2020). The HelD-RNAP complex was therefore proposed to be incompatible with active transcription. However, HelD in complex with RbpA and σ^A^ was recently shown to be involved in the initial steps of transcription initiation (Kovaľ *et al*., 2024). Although HelD function was mainly studied through *in vitro* transcription experiments, where the results depend on the HelD to RNAP ratio (Wiedermannová *et al*., 2014), the ratio of HelD to RNAP proteins in bacteria remains unknown. The HelD role *in vivo* is unclear and no systematic analysis of the effects of HelD on transcription *in vivo* has ever been performed.

Rifampicin, which inhibits bacterial RNA polymerase (Campbell *et al*, 2001), is currently used to treat tuberculosis. Rifampicin is often also recommended as therapy for various infections caused by nontuberculous mycobacteria, such as pulmonary disease caused by *Mycobacterium kansasii* or *Mycobacterium avium* complex (MAC) (Daley *et al*, 2020; Diel *et al*, 2018), although its effectiveness for the latter remains questionable (Schildkraut *et al*, 2023; van Ingen *et al*, 2024). The incidence of infections caused by non-tuberculous mycobacteria has been increasing globally (Johansen *et al*, 2020). Despite the increase, the factors contributing to rifampicin resistance in nontuberculous mycobacteria remain mostly unclear. *arr* gene encoding ADP-ribosyltransferase contributes to rifampicin resistance in *M. abscessus* (Morgado *et al*, 2021; Rominski *et al*, 2017) and *M. smegmatis* (Hetherington *et al*, 1995; Quan *et al*, 1999; Quan *et al*, 1997). *helD* was shown to confer rifampicin resistance in *M. abscessus* and *M. smegmatis* (Hurst-Hess *et al*, 2019; Hurst-Hess *et al*, 2021). The expression of HelD is induced by rifampicin and mediates rifampicin resistance in *M. smegmatis, M. abscessus* and *S. venezuelae* (Giddey *et al*, 2017; Hurst-Hess *et al*., 2019; Hurst-Hess *et al*., 2022; Surette *et al*., 2022). HelD in *M. smegmatis* was reported to improve growth in the presence of rifampicin in dilution spotting assays (Hurst-Hess *et al*., 2019; Hurst-Hess *et al*., 2022). In *S. venezuelae*, the primary function of HelD is proposed to be antibiotic resistance as HelD displace rifampicin from RNAP (Surette *et al*., 2022).

Here we analyzed the growth of major pathogenic nontuberculous mycobacteria in the presence of rifampicin. While the ability to growt in rifampicin partially correlated with the presence of *helD* in their genomes, *helD* was predominantly found in rapidly growing species, whereas slow-growing mycobacteria lacked *helD* homologs. We propose that the primary role of *helD* is to generally enhance transcription efficiency in rapidly growing mycobacteria. We demonstrate that HelD is present in substantial amounts during the growth of *Mycobacterium smegmatis* even in the absence of rifampicin and is located at the promoters of actively transcribed genes. We analyzed the transcriptome of a HelD knockout strain by RNA-seq and showed that HelD has a global effect on transcription. HelD enhances transcription during the exponential phase, a growth phase characterized by dense RNAP occupancy on highly expressed genes. During stationary phase of growth, HelD has the opposite effect on gene expression and reduces transcription. To explain this observation, we quantified the levels of HelD and major mycobacterial transcription factors (CarD, RbpA) and the intracellular GTP concentrations and detected notable differences during mycobacterial growth. Our results indicate that HelD is beneficial for mycobacteria not only to overcome rifampicin treatment but also to support rapid growth during exponential phase.

## RESULTS

### Nontuberculous mycobacteria differ significantly in their sensitivity to rifampicin

HelD has been shown to increase rifampicin resistance in *M. abscessus* (Hurst-Hess *et al*., 2022), but the data on rifampicin sensitivity in other mycobacterial species are limited or not available. To address this, we investigated rifampicin resistance in clinical isolates of mainly detected pathogenic mycobacterial species. Most isolates of *M. abscessus*, *M. avium*, and *Mycobacterium fortuitum* were able to grow at the highest rifampicin concentration used (50 μg/ml), whereas nearly all isolates of *Mycobacterium kansasii* were unable to grow at the lowest concentration (20 μg/ml) (Fig. 1).

**Figure 1.**
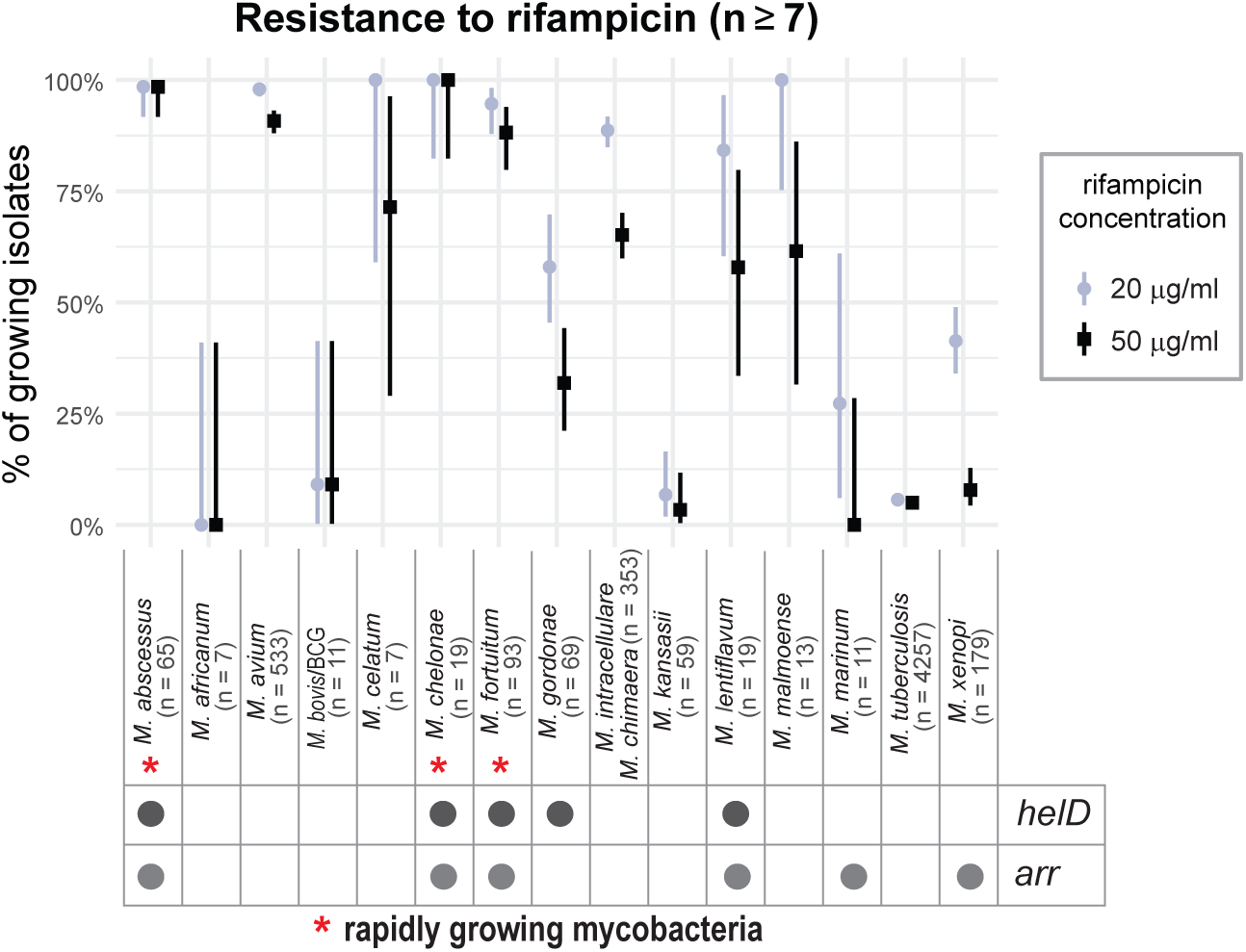
The presence of HelD improves the growth of cells treated by rifampicin. The analysis of rifampicin resistance in 15 mycobacterial species (a total of over 1,150 patient isolates), including three rapidly growing mycobacteria. Additionally, more than 3,600 *M. tuberculosis* clinical isolates were included for comparative analysis. The Y-axis shows the percentage of isolates for each species that were able to grow at a given concentration of rifampicin (50 μg/ml or 20 μg/ml). The presence of *helD* gene and *arr* gene (which encodes the ADP-ribosyltransferase) is indicated for each species by a dot at the bottom of the species table. The data provide relatively strong support for association between the presence of *helD* homologs and the observed resistance at the 50 μg/ml concentration (odds ratio: 33; 95% CI: 2.5 - 715; p = 0.01). For the 20 μg/ml concentration, the odds ratio is 18 (95% CI: 0.7 - 578; p = 0.07), meaning the data are consistent with an extremely strong positive association and a weak negative association. For *arr*, the remaining uncertainty is also very large and does not allow for any useful conclusions (all 95% CIs contain odds ratios 0.3 - 154, all p > 0.2). The conclusions are limited by the number of strains investigated, the lack of genotyping information at the sample level, and the resulting large statistical uncertainty.

The presence of *arr* and *helD* genes only partially explains the variability in rifampicin sensitivity among mycobacteria. Strikingly, we observed that *helD* was predominantly found in rapidly growing species, while slow-growing mycobacteria lacked *helD* homologs. The *helD* gene was present in all three rapidly growing species tested: *M. abscessus*, *M. fortuitum*, and *Mycobacterium chelonae* (Fig. 1). In contrast, many slow-growing pathogenic mycobacteria, such as *M. tuberculosis* or *M. avium*, lacked a *helD* homolog.

### HelD is found in the majority of rapidly growing mycobacteria

Mycobacteria are classified into two groups based on their growth rates: rapidly growing mycobacteria form visible colonies within seven days, whereas slow-growing mycobacteria require a longer period to form visible colonies. Most environmental, nonpathogenic, saprophytic mycobacteria belong to the rapidly growing group (Johansen *et al*., 2020). In contrast, the main pathogenic species belong to the slow-growing group, including the *M. tuberculosis* complex, *M. avium* complex (MAC), *M. kansasii*, *Mycobacterium xenopi*, and *Mycobacterium malmoense* (Fig. 1).

We found HelD homologs in almost 90 % of the investigated rapidly growing mycobacteria (Brown-Elliott & Philley, 2017) (Table 1), while most slow-growing mycobacteria lack HelD homologs (Bittner & Preheim, 2016) (Table 2). Thus, HelD is less common in slow-growing pathogenic species and is preferentially widespread among rapidly growing mycobacteria with transcriptionally highly active promoters and genes.

**Table 1.**
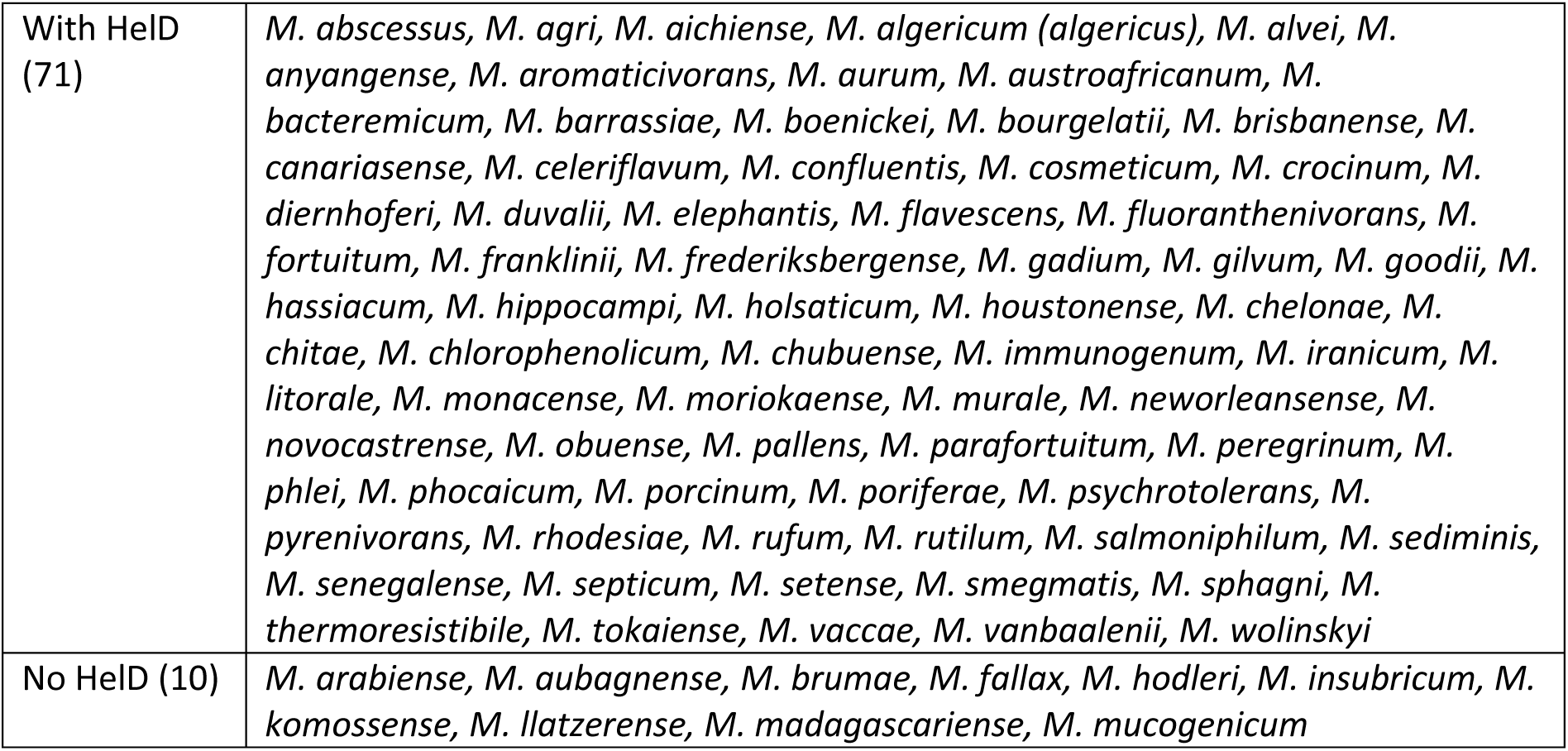
Rapidly growing mycobacteria with at least one HelD protein identified. (by BLASTp, OthoDB (Kuznetsov *et al*, 2023) or both).

**Table 2.**
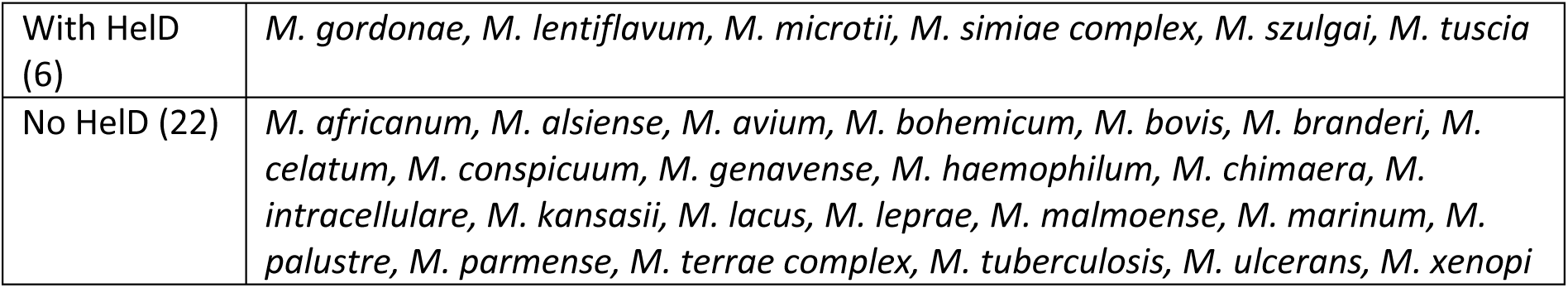
Slow-growing mycobacteria with at least one HelD protein identified. (by BLASTp, OthoDB (Kuznetsov *et al*., 2023) or both).

### HelD influences gene expression even in the absence of rifampicin

To investigate the role of HelD in fast-growing mycobacteria, we generated a *helD* deletion strain (Δ*helD*) in the rapidly growing *M. smegmatis*. In rich 7H9 medium, the growth of the Δ*helD* strain was not significantly different from that of wild type (wt) *M. smegmatis* (Fig. 2A). However, in the presence of rifampicin, the growth of the Δ*helD* was slower compared to the wt (Fig. 2A, 2B and Supplementary Fig. S1A). HelD immediately affected growth in liquid media with rifampicin, showing differences within two hours (Supplementary Fig. S1A). In addition, the strain overexpressing FLAG-tagged HelD showed improved growth in the presence of rifampicin compared to the wt (Supplementary Fig. S1B). The increased level of HelD protein even enabled *M. smegmatis* to grow at higher rifampicin concentrations.

**Figure 2.**
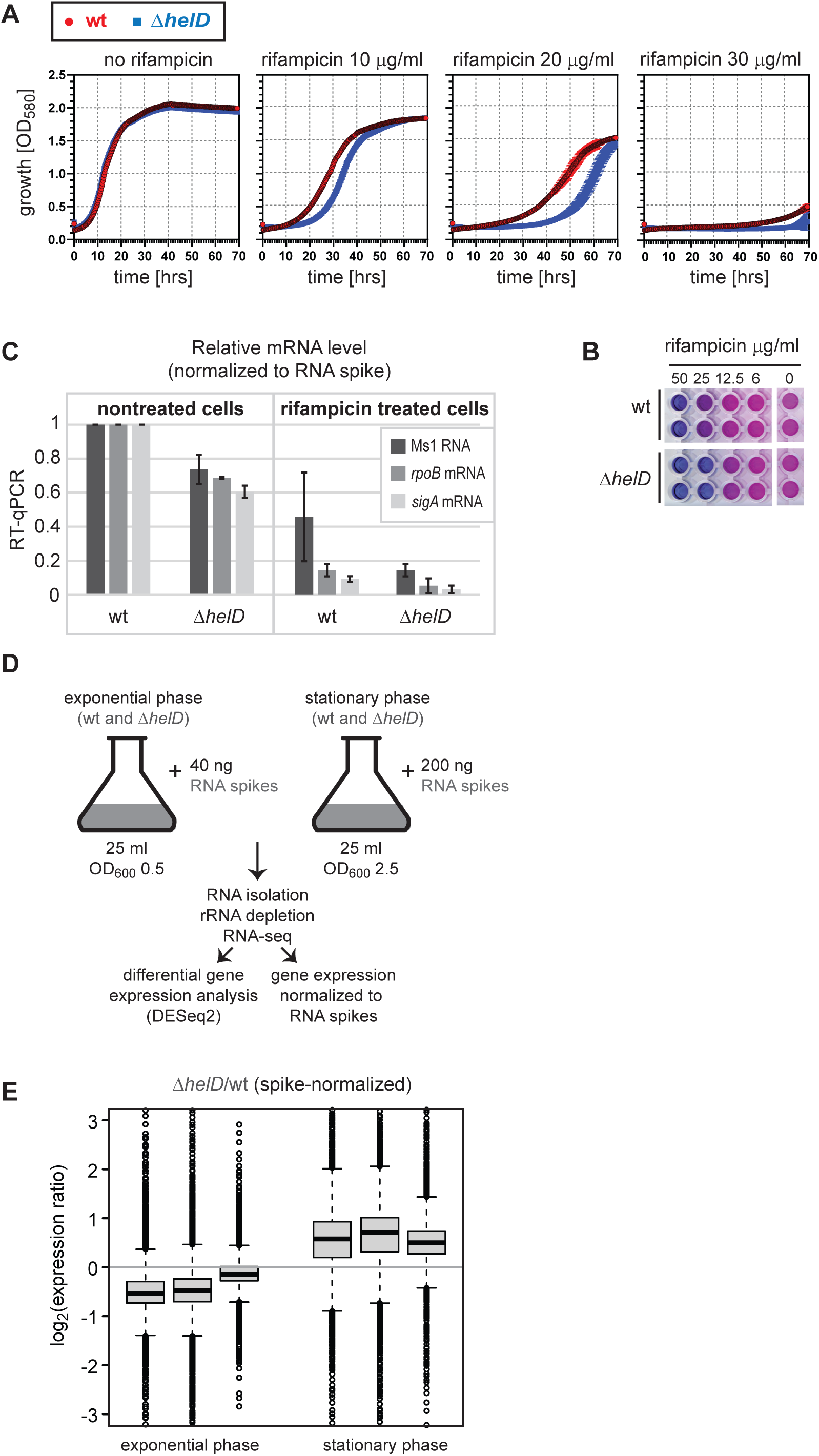
HelD has a global effect on gene expression in exponential and stationary phase. **A.** A growth curve of Δ*helD* and wt *M. smegmatis* strains in 7H9-ADC medium containing 0.5% (vol/vol) glycerol, 0.05% tyloxapol with and without rifampicin. **B.** Sensitivity to rifampicin was determined for both wt and Δ*helD M. smegmatis* strains by resazurin plate assay. Rifampicin sensitivity is higher for Δ*helD* [minimal inhibitory concentration (MIC) in between 12. 5 - 25 μg/ml] in comparison to wt strain (approximately 25 μg/ml). **C.** RT-qPCR analysis of gene expression of Ms1, *rpoB* and *sigA* RNAs in the Δ*helD* strain, normalized to the wt strain. Both strains were grown to exponential phase in the absence or presence of rifampicin (10 μg/ml), respectively. To accurately detect transcriptomic changes, we isolated RNA from equal numbers of cells and added four RNA spikes (eukaryotic RNA sequences of various lengths) at the start of RNA isolation. The mRNA levels were normalized to RNA spikes. **D.** The experimental design for RNA-seq of wt and Δ*helD* strains. Cells were harvested in exponential and stationary phase, respectively. RNA spikes were added in a quantity adjusted to the respective OD_600_ values, RNA was isolated, the rRNA was depleted and the RNA-seq was performed. To evaluate the effect of HelD deletion, both DESeq2 analysis and RNA spikes- based normalization were performed. **E.** The global gene expression changes in Δ*helD* strain compared to wt strain in exponential and stationary phase. Box plots show ratios of normalized expression of the Δ*helD* strain over the wt strain for each individual gene. Each box plot represents one biological replicate. The gene expressions were first normalized to RNA spikes and then the ratios were calculated.

HelD has been reported to restore transcription inhibited by rifampicin *in vitro* (Hurst-Hess *et al*., 2022; Kovaľ *et al*., 2024; Surette *et al*., 2022). In the presence of rifampicin, mRNA levels of three selected highly expressed genes (Ms1, *rpoB*, and *sigA*) were more reduced in Δ*helD* strain compared to the wt (Fig. 2C). HelD can therefore restore rifampicin-inhibited transcription *in vivo* as well.

Unexpectedly, in the exponentially growing control cells without rifampicin treatment, mRNA levels of Ms1, *rpoB*, and *sigA* also decreased in the Δ*helD* strain compared to the wt (Fig. 2C). HelD is therefore not only a rifampicin-specific transcription regulator but also regulates transcription under normal conditions.

### HelD is a global transcription regulator

To reveal the role of HelD in the regulation of gene expression in untreated cells, we performed RNA-seq using Δ*helD* and wt strain in both the exponential and stationary phases. RNA spikes were included to detect global transcriptomic changes (Fig. 2D). In the Δ*helD* strain, global gene expression decreased compared to wt in exponential phase. In stationary phase, global gene expression increased in Δ*helD* strain compared to wt (Fig. 2E). Therefore, HelD has opposite roles in transcription during exponential and stationary phases. While HelD increases RNA levels in the exponential phase, HelD has a negative effect on RNA levels in the stationary phase.

DESeq2 (which does not normalize to RNA spikes) detected that only two genes in exponential phase and nine genes in stationary phase were significantly differentially expressed in Δ*helD* strain compared to wt (Table 3 and Table 4, respectively), including *rpoC* gene encoding the RNAP β’ subunit. The RNAP β’ subunit mRNA level decreased in the Δ*helD* strain during the stationary phase, suggesting a reduced amount of RNAP and transcription; however, the global gene expression level increased in the Δ*helD* strain during this phase. Therefore, the specific gene expression changes detected by DESeq2 cannot explain the global shift in mRNA levels observed by RNA-seq.

**Table 3.** Differentially expressed genes in Δ*helD* strain compared to wt in exponential phase.

| Gene Name | Ratio $\Delta helD$/wt | function |
| --- | --- | --- |
| MSMEG_6359 | 0.28 | Protease, trypsin domain protein |
| MSMEG_1999 | 1.29 | hypothetical 47 aa long protein |

**Table 4.**
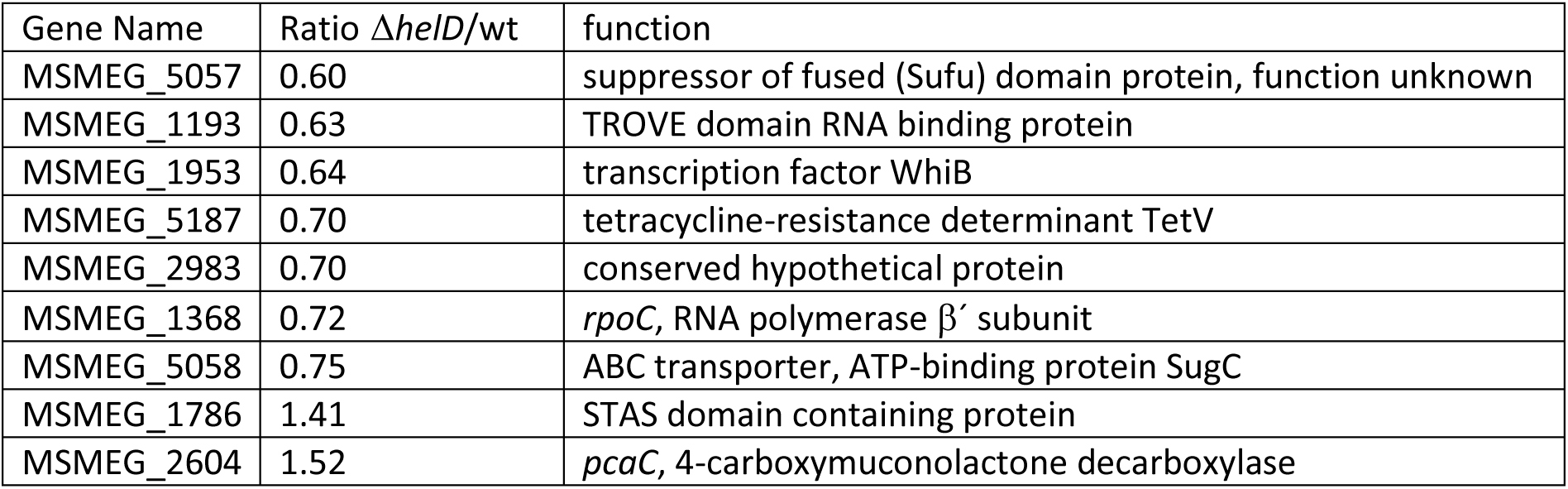
Differentially expressed genes in Δ*helD* strain compared to wt in stationary phase.

| Gene Name | Ratio $\Delta helD$/wt | function |
| --- | --- | --- |
| MSMEG_5057 | 0.60 | suppressor of fused (Sufu) domain protein, function unknown |
| MSMEG_1193 | 0.63 | TROVE domain RNA binding protein |
| MSMEG_1953 | 0.64 | transcription factor WhiB |
| MSMEG_5187 | 0.70 | tetracycline-resistance determinant TetV |
| MSMEG_2983 | 0.70 | conserved hypothetical protein |
| MSMEG_1368 | 0.72 | *rpoC*, RNA polymerase $\beta'$ subunit |
| MSMEG_5058 | 0.75 | ABC transporter, ATP-binding protein SugC |
| MSMEG_1786 | 1.41 | STAS domain containing protein |
| MSMEG_2604 | 1.52 | *pcaC*, 4-carboxymuconolactone decarboxylase |

### HelD associates with RNAP involved in transcription *in vivo*

If HelD is involved in transcription regulation, we could detect it directly on the genomic DNA through chromatin immunoprecipitation (ChIP). To identify specific genes associated with HelD by ChIP-seq (Fig. 3A) we used a strain overexpressing HelD-FLAG (with a HelD- FLAG to RNAP ratio of approximately 1:1, Supplementary Fig. S2), increasing the likelihood of observing also transient HelD-RNAP-DNA complexes.

**Figure 3.**
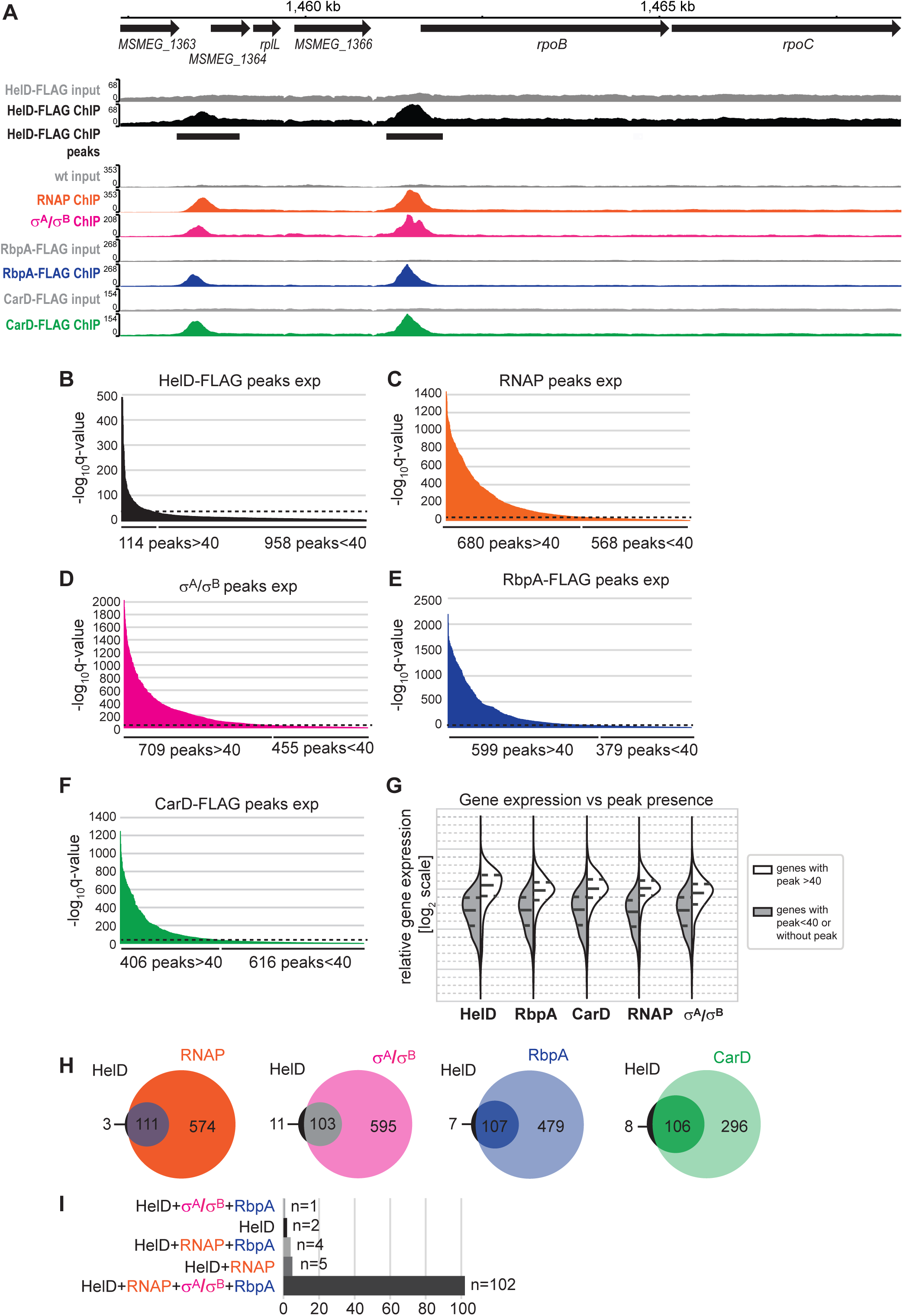
HelD-FLAG distribution overlaps with CarD-FLAG, RbpA-FLAG, RNAP and σ^A^/σ^B^. **A.** Exponential phase ChIP-seq profiles of HelD-FLAG, RNAP, σ^A^/σ^B^, RbpA-FLAG and CarD-FLAG in a representative region of *M. smegmatis* genome which includes *rpoB-rpoC* operon. Statistically significant HelD-FLAG peaks (-log_10_q-value>40) are labelled by boxes. The HelD-FLAG peaks were detected at promoters and overlap with RNAP, σ^A^/σ^B^, RbpA- FLAG and CarD-FLAG peaks. This panel was adopted from the msmegseq.elixir-czech.cz webpage. Exponential phase ChIP-seq peaks detected for HelD-FLAG **(B)**, RNAP **(C)**, σ^A^/σ^B^ **(D)**, RbpA-FLAG **(E)** and CarD-FLAG **(F)**. The y-axis shows the statistical significance of each individual peak, peaks with -log_10_q-value>40 were used for further analysis. **G.** The significant peaks (-log_10_q-value>40, shown in red on the right in violin plots) are present on genes with higher expression levels. Genes without the detected peaks or with less significant peaks (-log_10_q-value<40 are shown in grey on the left in violin plots) are less expressed. Dashed lines in violin plots represent the lower and upper quartiles. The y-axis shows the gene expression in exponential phase from the previously published *M. smegmatis* RNA-seq dataset (Šiková *et al*., 2019). **H.** Venn diagrams showing the overlap of ChIP-seq peaks (-log_10_q- value>40). Only significant peaks with -log_10_q-value>40 were considered. **I.** Out of 114 HelD- FLAG peaks (-log_10_q-value>40), 102 peaks overlap with RNAP, σ^A^/σ^B^ and RbpA-FLAG peaks (-log_10_q-value>40). Only two HelD-FLAG do not overlap with RNAP, σ^A^/σ^B^ and RbpA-FLAG peaks.

With ChIP-seq, we detected 114 peaks with -log_10_q-value>40 which were used for further analysis (for all HelD-FLAG ChIP-seq peaks, see Supplementary Table S1). An example of ChIP-seq data for ∼11kb genomic region including *rpoB-rpoC* promoter is shown in Fig. 3A. *rpoB* and *rpoC* encode the largest subunits of RNAP β and β′, respectively, and HelD-FLAG binds to their promoter. HelD was mainly bound to the highly expressed genes including both ribosomal rRNA operons, tRNAs, genes encoding RNAP subunits (β, β′, α, ω) or ribosomal proteins (Supplementary Table S1).

Then, we compared HelD-bound genomic regions with the binding sites of RNAP, σ^A^, RbpA and CarD. ChIP-seq was performed previously for CarD (Landick *et al*, 2014) showing that CarD binds promoters that are also occupied by RNAP and σ^A^. No ChIP data is available for RbpA in *M. smegmatis*.

We performed ChIP-seq in strains expressing RbpA-FLAG and CarD-FLAG. FLAG-tagged strains are well established for mycobacterial ChIP-seq (Minch *et al*, 2015) and enable us to compare different transcription factors with the same antibody. For an unknown reason, FLAG-tagged σ^A^ cannot be efficiently used for chromatin immunoprecipitations, as observed also by others (Hurst-Hess *et al*., 2019). We performed σ^A^ ChIP with an anti-σ^A^ antibody recently in the wt strain (Vanková Hausnerová *et al*, 2024), which is the parental strain for all FLAG-tagged strains used in this study. The anti-σ^A^ antibody also recognizes σ^B^, which has a very similar amino acid sequence in *M. smegmatis* (Singha *et al*, 2023; Vanková Hausnerová *et al*., 2024). Hence, we will refer to this antibody as anti-σ^A^/σ^B^. RNAP ChIP-seq was also performed recently using anti-RNAP antibody (Vanková Hausnerová *et al*., 2024).

We detected considerably more peaks with higher statistical significance (-log_10_q-value > 40) for RNAP, σ^A^/σ^B^, RbpA, and CarD than for HelD (Fig. 3B versus Fig. 3C, D, E and F), indicating that the association of HelD with the transcription machinery might be transient or less frequent. HelD, like RNAP, σ^A^/σ^B^, RbpA, and CarD, is associated with genes that are highly expressed (Fig. 3G), which suggests that HelD is present at actively transcribed genes. All ChIP-seq data and detected peaks are available at the msmegseq.elixir-czech.cz webpage with the integrated IGV genome browser (Robinson *et al*, 2023).

Almost all HelD peaks overlapped with RNAP, σ^A^/σ^B^, RbpA, or CarD (Fig. 3H). Additionally, there were only two genes where HelD was the sole binding factor (Fig. 3I). In conclusion, HelD associates with the same genomic regions as RNAP, σ^A^/σ^B^, CarD and RbpA and it binds to the promoters *in vivo*.

The ChIP-seq data showed that RNAP peaks and σ^A^/σ^B^ peaks tend to be higher when they overlap with HelD peaks (Fig. 4A and B). This correlation indicates that HelD is recruited to DNA by RNAP and/or σ^A^/σ^B^, which is consistent with the recently revealed RNAP-σ^A^- RbpA-HelD-DNA cryo-EM structure (Kovaľ *et al*., 2024). The presence of HelD correlates with increased gene expression also on a subset of tRNA genes. tRNA genes associated with HelD had higher expression level than those not associated with HelD (Fig. 4C and D). Based on ChIP-seq data, HelD is involved in transcription initiation as well, not only in the release of stalled RNAPs within genes or in transcription termination (Kouba *et al*., 2020) and both processes might be coupled (Kovaľ *et al*., 2024).

**Figure 4.**
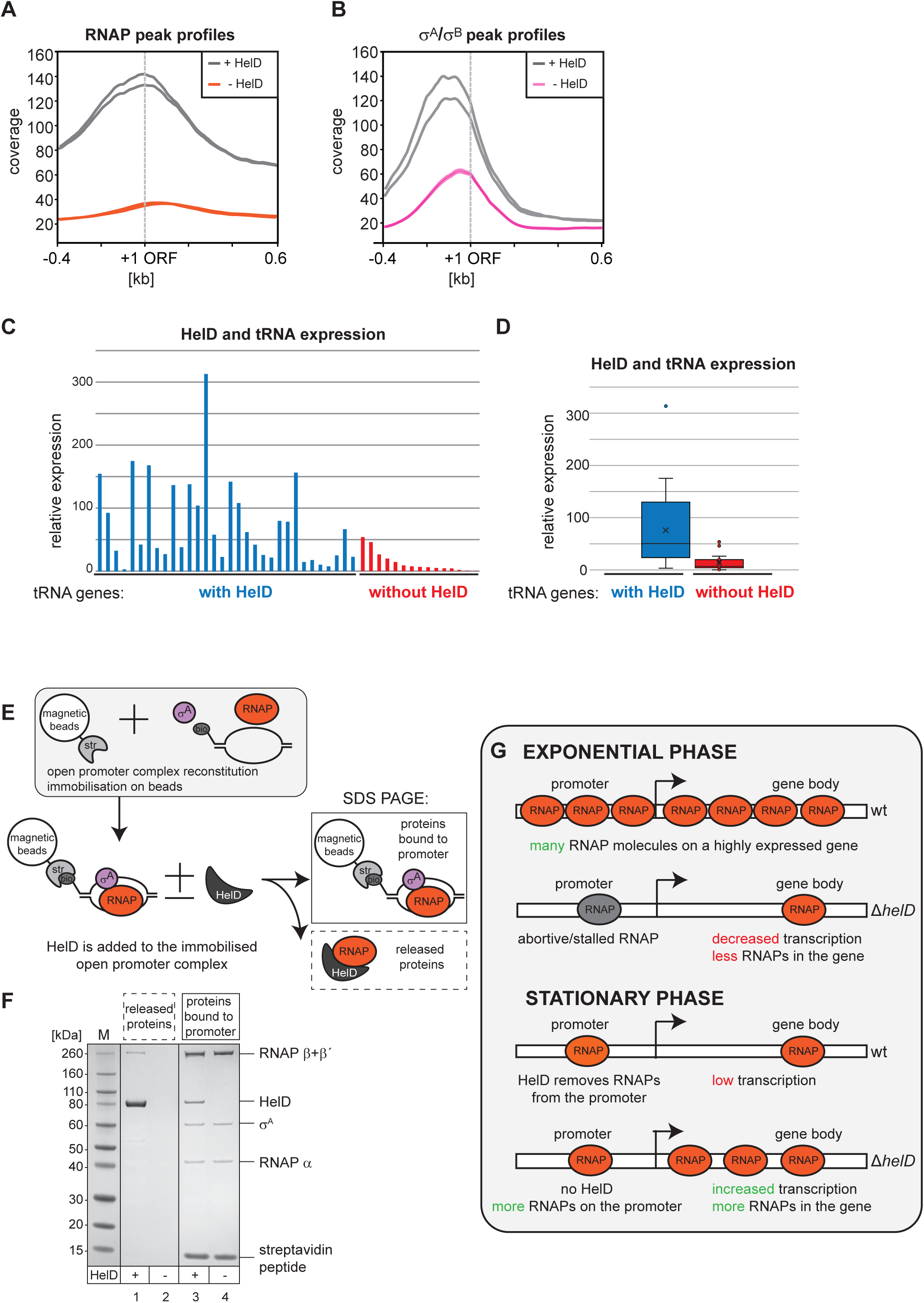
HelD is localized on promoters of highly expressed genes. **A, B.** ChIP-seq coverage profiles (from -0.4 kb to +0.6 kb around ORF start site). In **A**, grey lines show RNAP peaks that overlap with HelD-FLAG peaks, while orange lines show RNAP peaks that do not overlap with HelD-FLAG peaks. In **B**, grey lines show σ^A^/σ^B^ peaks that overlap with HelD-FLAG peaks, while pink lines show σ^A^/σ^B^ peaks that do not overlap with HelD-FLAG peaks. Each line represents an individual biological replicate. Note that the RNAP and σ^A^/σ^B^ peaks overlapping with HelD-FLAG have higher coverage compared to RNAP and σ^A^/σ^B^ peaks that do not overlap with HelD-FLAG. **C, D.** In exponential phase ChIP-seq, HelD- FLAG peak (-log_10_q-value>40) was detected at 32 tRNA gene promoters (in blue) and not detected at 15 tRNA promoters (in red). The relative expression in exponential phase for each tRNA is shown in **C**, HelD-occupied tRNA genes are sorted according to the significance (-log_10_q-value) of the detected HelD peaks. **D.** A box plot showing exponential phase relative gene expression of tRNA genes occupied by HelD (in blue) or devoid of HelD (in red). The relative tRNA expression levels are adopted from RNA-seq dataset that was published previously (Šiková *et al*., 2019). **E.** The experimental scheme of the *in vitro* RNAP release assay. The RNAP open promoter complex consisting of two annealed DNA oligos forming a bubble, RNAP and σ^A^ proteins was reconstituted and bound to beads via streptavidin-biotin interaction. The complex was incubated in the presence or absence of HelD protein, and subsequently, the proteins that remained in the complex (were bound to the beads) and the released proteins (present in the supernatant) were visualized using SDS-PAGE. **F.** The released proteins and those that remained bound to the open promoter complex were resolved on the SDS-PAGE. Lines 1 and 3 show the assay in which HelD was present and lines 2 and 4 show the assay in which HelD was absent. The released RNAP was detected only in the presence of HelD (line 1). **G.** A model of HelD function in exponential and stationary phase. In exponential phase, when abortive RNAP stalls at promoters, transcription cannot proceed, leading to a decrease in mRNA levels and the reduction of transcribing RNAP along genes. HelD, known for its ability to release stalled RNAP, acts at promoters, facilitating the release of abortive/stalled RNAP and allowing access for other RNAPs, thereby enhancing transcription in the exponential phase. In stationary phase, diminished levels of RbpA and CarD slow down transcription initiation. In the wt strain, HelD-mediated RNAP release may not only target abortive/stalled RNAPs but also those awaiting transcription initiation at promoters. In the Δ*helD* strain, these RNAPs remain at the promoters, leading to an increased transcription.

### HelD can release RNAP from transcription initiation complexes

HelD, which is located at promoters (Fig. 3A), might release abortive RNAPs (RNAPs that synthesize short RNA fragments but fail to enter productive elongation (Chen *et al*, 2021)) from the promoter and make the promoter accessible for other RNAPs which could increase transcription in exponential phase in *M. smegmatis*. In the exponential phase, HelD is associated with highly expressed (Fig. 3G), RNAP-occupied genes (Fig. 4A) and HelD presence increases gene expression (Fig. 2E and 4C).

We assembled an open promoter complex including RNAP with σ^A^ and biotinylated promoter DNA (Hubin *et al*, 2017b; Kovaľ *et al*., 2024) on streptavidin beads (Fig. 4E, Supplementary Fig. S3). When we added HelD to this complex, HelD was able to release a fraction of RNAP (Fig. 4F). Therefore, HelD is capable of disassembling transcription initiation complexes. HelD can probably release any abortive RNAP complexes and directly recycle them into transcription initiation (Kouba *et al*., 2020; Kovaľ *et al*., 2024).

In exponential phase, highly expressed genes are occupied by large amounts of RNAP due to the high level of transcription (Fig. 4G). We propose that when abortive RNAP complexes cannot be released or recycled from the promoter in the Δ*helD* strain, transcription cannot proceed, leading to a reduction in both mRNA level and the amount of transcribing RNAPs along the gene. Consistent with this, by ChIP we detected a lower RNAP level at the 16S rRNA gene in the Δ*helD* strain compared to the wt (Supplementary Fig. S4A).

In contrast, during the stationary phase, the presence of HelD decreases gene expression (Fig. 2E). Transcription initiation may generally be less frequent and efficient, and HelD releases not only abortive/stalled RNAPs but also RNAPs at promoters that have not yet initiated transcription. These RNAPs are not released from promoters in the *ΔhelD* strain, leading to an accumulation of RNAPs not only within genes but also at promoters (Fig. 4G). In the Δ*helD* strain, RNAP levels in stationary phase increased at the 16S rRNA gene, Ms1 gene, and *rpoB-rpoC* promoter compared to the wt as quantified by ChIP (Supplementary Fig. S4B and S4C) which supports our model.

### GTP concentration decreases in stationary phase *M. smegmatis*

Despite the widespread use of *in vitro* methods to study mycobacterial transcription, almost no data are available on the *in vivo* concentrations of basic transcription factors in mycobacteria during the exponential and stationary phases. To fully explain HelD’s opposing effects on transcription during these phases, we measured differences in the availability of transcription factors and initiation nucleotides.

Transcription initiation is regulated by the level of initiation nucleoside triphosphates (iNTPs) (Krásný *et al*, 2008). In ∼55 % of all *M. smegmatis* transcripts expressed in exponential and stationary phase, G was present at the +1 position (Fig. 5A, the top logos) (Šiková *et al*, 2019). Transcription of both highly and lowly expressed genes initiates mainly with GTP and can be affected by GTP concentration in *M. smegmatis*. Therefore, we measured the intracellular levels of GTP in exponential and stationary phase cells (Fig. 5B). The GTP level decreased approximately three times in stationary phase when compared to exponential phase indicating lower and less frequent transcription initiation in stationary phase.

**Figure 5.**
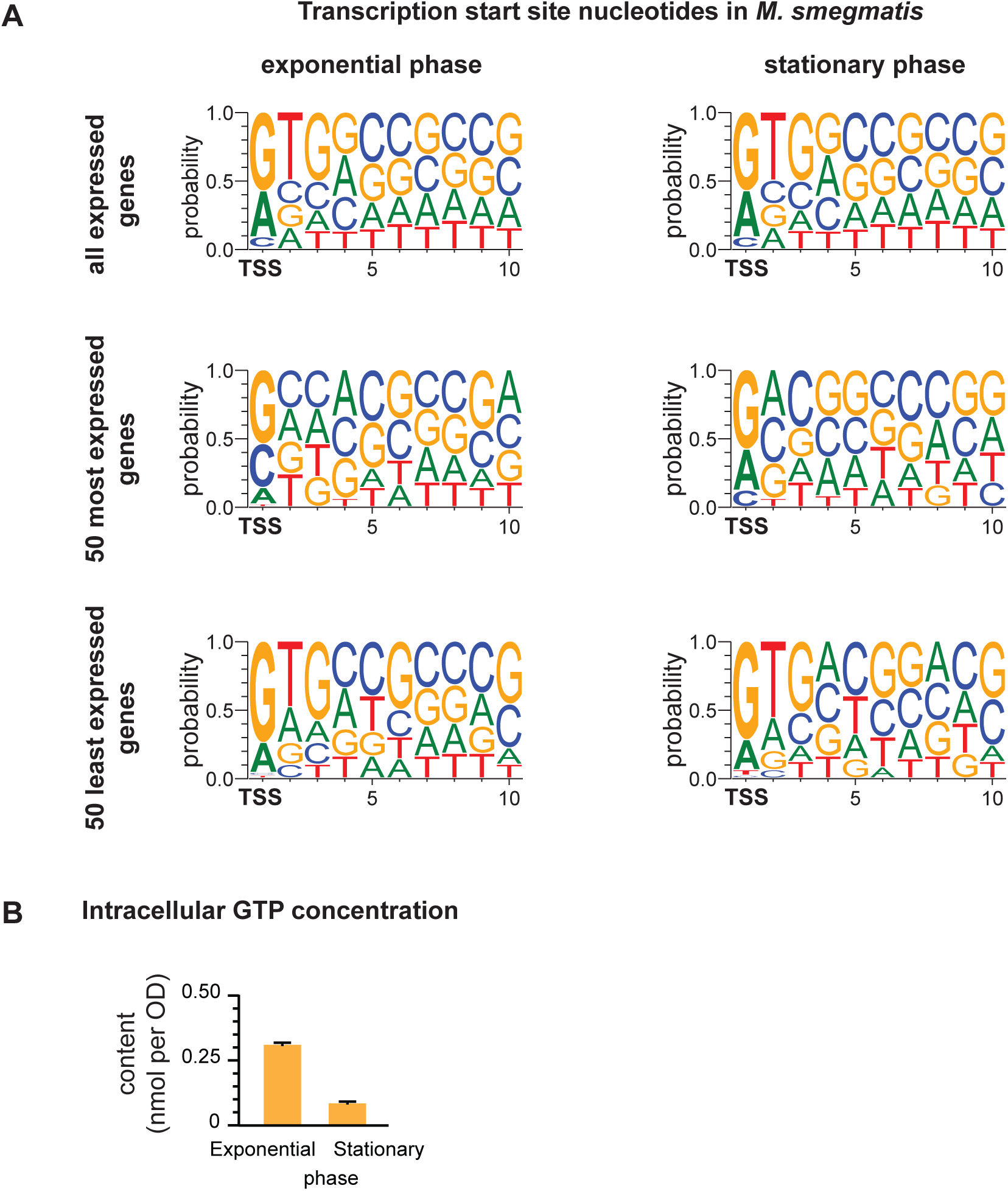
*M. smegmatis* transcripts are characterized by a high proportion of G on +1 position. **A.** Logos representing a sequence of the first 10 nucleotides of all expressed genes, 50 most expressed genes and 50 least expressed genes in exponential and stationary phase, respectively. The transcription start sites (TSS) are highlighted in bold. The related gene expression data were published previously (Šiková *et al*., 2019). The top 50 genes with the highest expression were characterized by a similar frequency of +1 G (Fig. 5A, middle logos), while the 50 least expressed genes had even higher frequency of +1 G (∼75 %, see Fig. 5A, bottom logos). **B.** The intracellular levels of guanosine triphosphate (GTP) in exponential and stationary phase.

### *M. smegmatis* has more RNAP than HelD molecules

To determine *in vivo* concentrations of individual components of the mycobacterial transcription machinery, we produced custom antibodies for CarD, RbpA, and HelD. We focused on the protein levels, as CarD mRNA levels do not correlate with CarD protein levels due to extensive post-transcriptional regulation (Li *et al*, 2022). The sonication protocol to prepare exponential and stationary phase *M. smegmatis* lysates was optimized, ensuring that over 90 % of proteins were present in the lysates (Supplementary Fig. S5).

First, we determined the absolute amounts of endogenous HelD and RNAP in exponential and stationary phase lysates by comparing them with varying quantities of *in vitro* purified HelD and RNAP using western blotting (Fig. 6A, lanes “wt”, and 6B). One µg of exponential phase lysate from the wt strain contained 115 fmol of RNAP and 18 fmol of HelD, resulting in an RNAP:HelD ratio of approximately 6:1. In stationary phase, the RNAP:HelD ratio was around 4:1. The HelD-FLAG overexpression strain was used as a positive control for HelD antibody and confirmed that the ratio of HelD to RNAP was approximately 1:1 in the HelD-FLAG ChIP-seq from the exponential phase (Fig. 6A, lanes “FLAG”).

**Figure 6.**
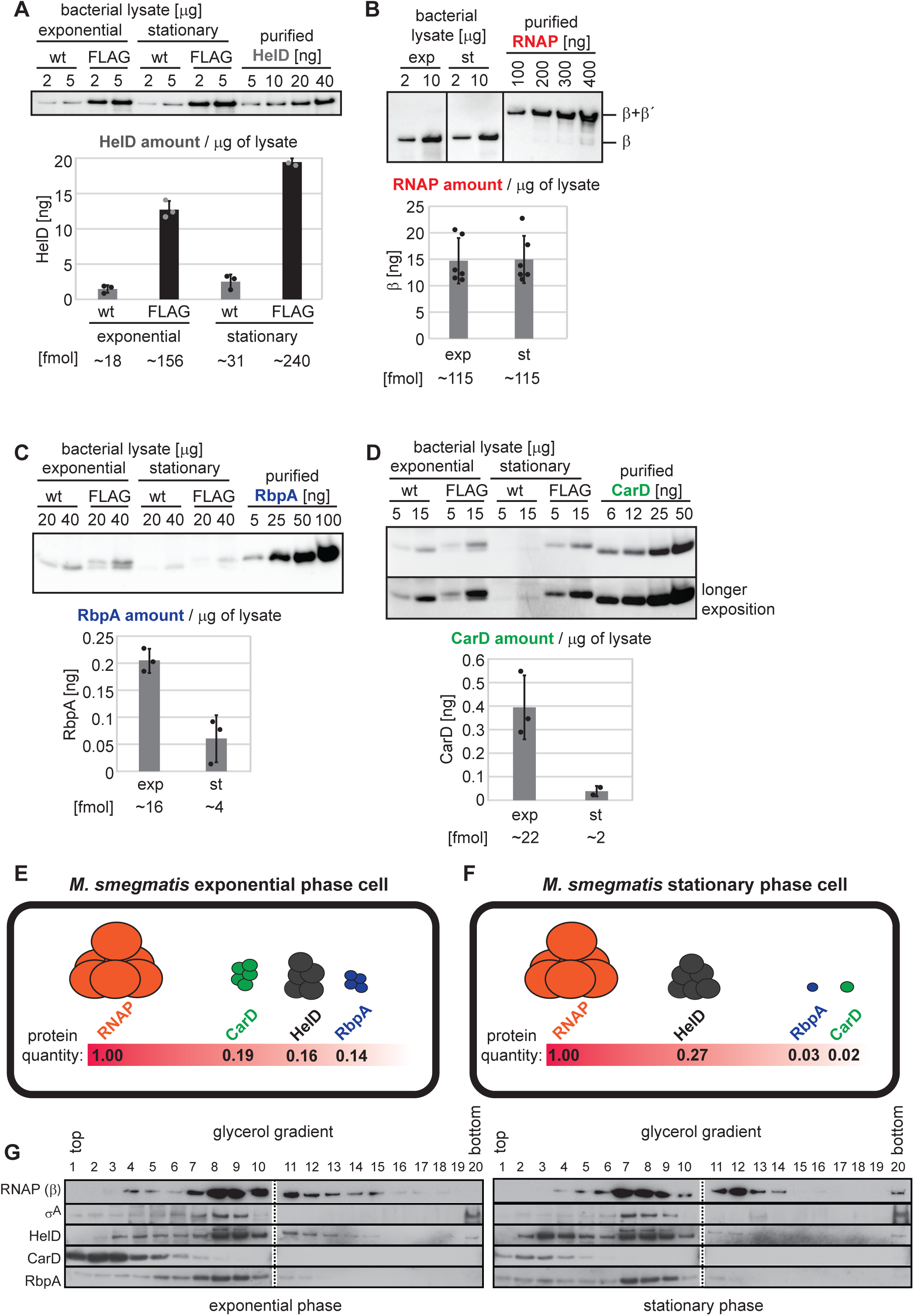
HelD amount *in vivo* relative to RNAP, CarD and RbpA. **A, B, C, D.** The amounts of individual proteins per 1 μg of exponential and stationary phase lysates were quantified by western blotting using *in vitro* purified proteins to generate standard curves for calculations. In addition, lysates with overexpressed ("FLAG") FLAG-tagged HelD (**A**), RbpA (**C**) and CarD (**D**) were used to determine the protein levels after overexpression. To detect HelD, RbpA and CarD, we produced our own mouse anti-HelD, anti-RbpA and anti-CarD antibodies, the amount of RNAP was quantified with a commercially available anti-RNAP β antibody (8RB13). The amount of HelD increased approximately eight times in HelD-FLAG strain in exponential phase when compared to wt (**A**). *In vitro* purified RNAP is composed of β and β′ subunits connected by a linker (Kouba *et al*., 2019), which was also considered when the molar concentration was calculated (**B**). **E. F.** The approximate number of molecules of CarD, RbpA and HelD per one molecule of RNAP in exponential phase **(E)** and stationary phase **(F)**. **G.** The complexes from *M. smegmatis* exponential and stationary phase lysates were fractionated by glycerol gradient ultracentrifugation. The presence of RNAP, σ^A^, HelD, RbpA and CarD in individual gradient fractions was determined by western blotting. While CarD was mainly detected at the top of the gradient, HelD and RbpA were mainly detected in the same fractions as RNAP.

### In stationary phase, CarD and RbpA levels decreased, while HelD increased

While the level of RNAP remained constant in both exponential and stationary phases (Fig. 6B), and the level of HelD partially increased in stationary phase (Fig. 6A), we observed a ∼ten-fold decrease in CarD, as previously reported (Li *et al*., 2022) (Fig. 6C). Notably, for the first time, we also observed a ∼four-fold decrease in RbpA during stationary phase (Fig. 6D). In exponential phase, the ratios of HelD, RbpA, and CarD to RNAP were similar. However, in the stationary phase, CarD and RbpA were significantly less abundant relative to RNAP (Fig. 6E and F).

In addition, not all CarD was bound to RNAP. To determine the proportion of CarD that was associated with RNAP, we fractionated lysates from exponential and stationary phase *M. smegmatis* by ultracentrifugation in glycerol gradients (Hnilicova *et al*, 2014). While RNAP complex mainly sedimented in fractions 7 to 13, CarD sedimented in fractions 1 to 6 (Fig. 6G). This indicates that the majority of CarD is not in complex with RNAP. In contrast, the majority of HelD co-sedimented with RNAP during the exponential phase (Fig. 6G, fractions 8 to 10). Similarly, most RbpA co-sedimented with RNAP, suggesting that RbpA is almost entirely bound to RNAP, with little to no free RbpA present in the cell (Fig. 6G).

We were unable to detect CarD and HelD in the same complex *in vivo*. When CarD-FLAG was immunoprecipitated, HelD was almost not detected among the co-immunoprecipitated proteins, neither in exponential phase (Fig. 7A, lane 1) nor in stationary phase (Fig. 7A, lane 3). When RbpA-FLAG was immunoprecipitated, HelD was detected in both exponential and stationary phase (Fig. 7A, lanes 2 and 4). CarD and HelD can associate with the same promoters (Fig. 3H), but not within the same complexes, which is consistent with the recently published HelD-FLAG interactome where CarD is absent and which is also supported by cryo-EM structural data (Kovaľ *et al*., 2024).

**Figure 7.**
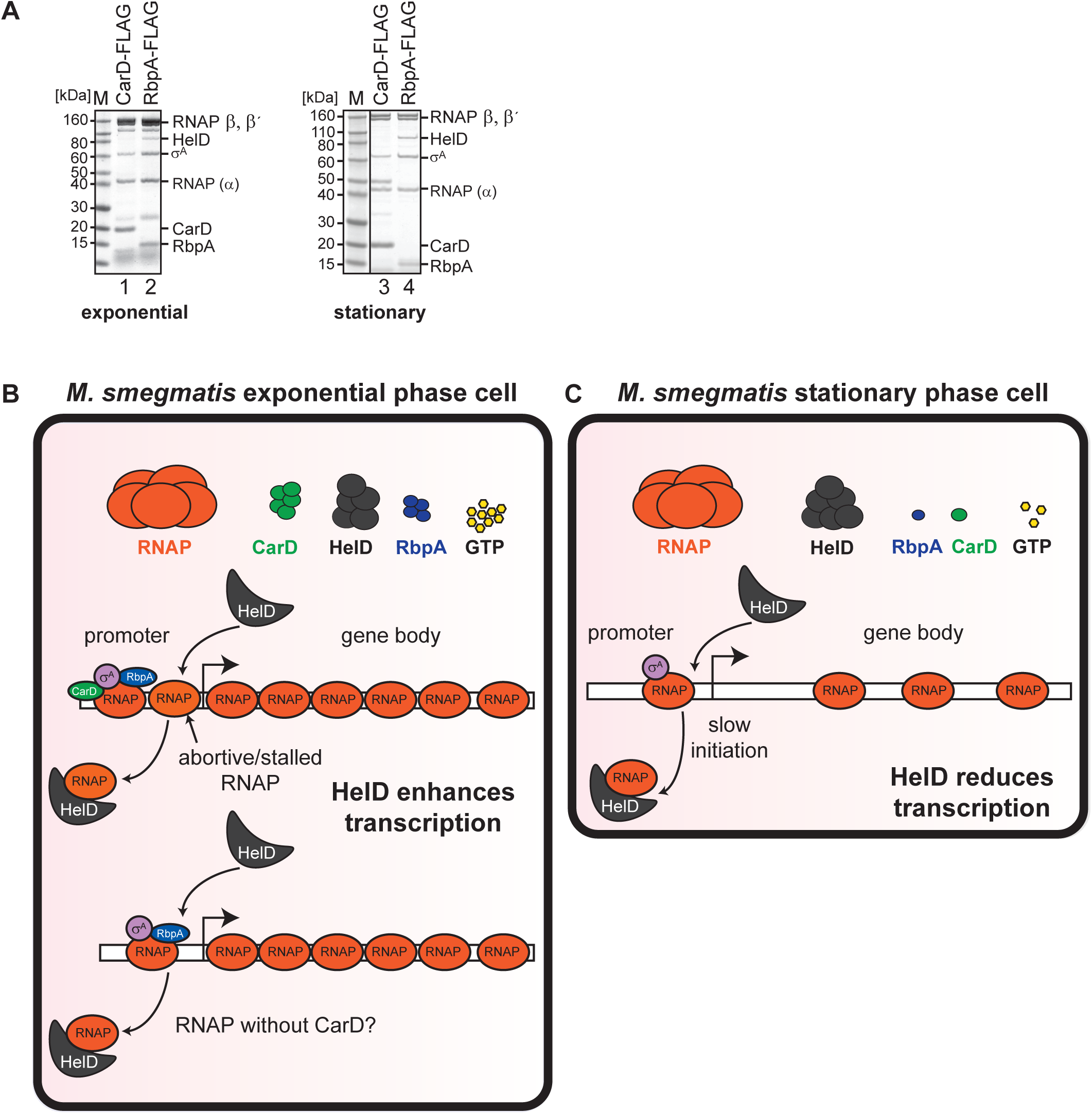
Molecular model of the mechanism of *M. smegmatis* HelD action *in vivo.* **A.** CarD-FLAG and RpbA-FLAG were pulled down from exponential and stationary phase cells and co-immunoprecipitated proteins were resolved by SDS-PAGE. The identity of protein bands was determined by mass spectrometry. **B. C.** We propose that in exponential phase (**B**), HelD is involved in removing abortive/stalled RNAPs or CarD-less RNAPs from the promoters of highly expressed genes. This results in the overall increase of transcription in exponential phase when HelD is present. In stationary phase (**C**), HelD targets not only abortive/stalled RNAPs but also those waiting for CarD, RbpA, or GTP. This leads to the overall reduction of transcription in stationary phase when HelD is present.

Our data suggest that the levels of CarD and RbpA in the mycobacterial cell are lower compared to RNAP, particularly during the stationary phase (Fig. 6F). Moreover, only a small fraction of the total CarD molecules is bound to RNAP during both exponential and stationary phases (Fig. 6G). Therefore, although both CarD and RbpA proteins bind RNAP (Hu *et al*., 2012; Stallings *et al*., 2009; Weiss *et al*, 2012), CarD appears to associate with RNAP only transiently and acts as the limiting transcription regulator. This highlights that bacterial transcription regulation is a dynamic process and underscores the importance of *in vivo* data for complementing *in vitro*-based models.

We demonstrate that HelD associates with promoters and can release RNAP from promoters both *in vivo* and *in vitro*. The role of the HelD protein differs between the exponential and stationary phases, which we explain by the varying concentrations of the transcription factors CarD and RbpA, as well as the primary initiation nucleotide, GTP. HelD is not only a gene implicated in rifampicin resistance; rather, in rapidly growing mycobacteria, it functions as a general transcription factor capable of recycling RNAP.

## DISCUSSION

Rifampicin treatment induces an increase in HelD expression (Giddey *et al*., 2017; Hurst-Hess *et al*., 2019), but in *M. smegmatis*, a substantial level of HelD is present even without rifampicin treatment. This suggests that HelD has additional roles beyond protecting RNAP from rifampicin.

*B. subtilis* HelD is believed to promote RNAP dimerization as (RNAP-HelD)2 dimers have been observed (Newing *et al*., 2020), and it is speculated that this form of RNAP is in a hibernating and non-transcribing state. *M. smegmatis* HelD was also proposed to store RNAP in an inactive state (Kouba *et al*., 2020). To answer whether RNAP molecules could be trapped by HelD in *M. smegmatis*, we measured the amounts of HelD and RNAP *in vivo*. Based on HelD:RNAP ratio, HelD does not have the capacity to sequester more than 15 % of RNAPs in exponential phase and 25 % in stationary phase. Given the excess of RNAP over HelD, we concluded that HelD is unlikely to serve as a major storage factor for inactive RNAP molecules. Importantly, overexpression of HelD did not negatively affect growth of the strain (Supplementary Fig. S1B), indicating that, at least during exponential phase, HelD does not significantly sequester and inactivate RNAP in *M. smegmatis*.

Here, we propose direct involvement of HelD in global transcription regulation. We observed that HelD interacts with promoters (Fig. 3A), which is consistent with the presence of RbpA, and the primary sigma factor, σ^A^ in complex with HelD (Kouba *et al*., 2020). Recently, the *in vitro* cryo-EM structure of RNAP-σ^A^-RbpA-HelD-DNA transcription initiation complex was revealed (Kovaľ *et al*., 2024), showing that HelD can associate with DNA promoters, which is in agreement with our ChIP-seq *in vivo* data. HelD ChIP-seq signal overlaps with RNAP and σ^A^/σ^B^ distribution along the *M. smegmatis* genome and HelD binds to a subset of promoters that can also associate with CarD and RbpA.

Mycobacterial HelD has been previously suggested to help recycle stalled RNAP molecules (Kouba *et al*., 2020; Kovaľ *et al*., 2024; Sudzinová *et al*, 2022). *M. abscessus* HelD was shown to partially disrupt the interaction between RNAP and promoter DNA, but σ^A^ was absent in the tested RNAP-DNA complex (Hurst-Hess *et al*., 2022). Based on cryo-EM data, a specific binding site formed by σ^A^/σ^B^ for HelD in the context of initiating RNAP was identified (Kovaľ *et al*., 2024) and our data show that HelD can release RNAP from RNAP-σ^A^-DNA open promoter complex both *in vitro* (Fig. 4F) and *in vivo* (Supplementary Fig. S4C).

In exponential phase, HelD in complex with RNAP mainly associates with promoters of highly expressed genes, such as the majority of tRNA genes (Fig. 4C) and generally increases gene expression. We hypothesize that this can be explained by a high number of RNAP molecules moving along these busy promoters. During exponential growth, there is a high demand for tRNA production, and HelD probably helps to keep these promoters fully accessible for the numerous RNAP molecules, and this in turn increases gene expression (Fig. 4G). It is worth mentioning that *helD* gene is absent from many slow-growing mycobacteria and present in many rapidly growing mycobacteria (including *M. smegmatis*) which correlates with a higher transcription rate in these species.

In contrast to exponential phase, stationary phase cells produce approximately 10 % more mRNA in the absence of HelD. In order to explain this observation, we need to take into account two factors that differ between exponential and stationary phase *M. smegmatis* cells: i) the level of CarD and RbpA proteins drops in stationary phase (there are 11 times fewer CarD molecules and four times fewer RbpA molecules in stationary phase compared to exponential phase (Fig. 6), and ii) the GTP content of stationary phase cells is approximately three times lower than in exponential phase cells (Fig. 5B). In addition, mycobacterial RNAP is also highly associated with Ms1 regulatory RNA during stationary phase (Hnilicova *et al*., 2014; Vanková Hausnerová *et al*., 2024; Vaňková Hausnerová *et al*, 2022).

Mycobacterial transcription is characterized by an unstable open promoter complex (RP_o_) (Davis *et al*, 2015; Garner *et al*, 2017; Jensen *et al*, 2019). To efficiently transition from the initiation to elongation, mycobacteria require the transcription factors, CarD and RbpA (Bortoluzzi *et al*, 2013; Hu *et al*., 2012; Hubin *et al*, 2015; Srivastava *et al*., 2013; Stallings *et al*., 2009). CarD has been demonstrated to stabilize the mycobacterial RP_o_ in a concentration-dependent manner – the higher the concentration of CarD, the more stable the RP_o_ (Rammohan *et al*., 2015). We hypothesize that in stationary phase, the decreased level of CarD and RbpA, reflects the general decline of transcription and thus lower demand for transcription factors and eventually results in a slow-down of RP_o_ formation kinetics (Jensen *et al*., 2019; Rammohan *et al*., 2015). Subsequently, the RNAPs could thus be more susceptible to HelD-mediated release of RNAP from the promoter (Fig. 7C). We performed RNAP ChIP in Δ*helD* strain and wt strain to demonstrate HelD’s effect on RNAP genome association *in vivo* on three highly expressed genes. In stationary phase, we were able to detect more RNAP molecules at *rpoB* promoter in Δ*helD* than in wt (Supplementary Fig. S4C). This finding is consistent with the increased global gene expression in Δ*helD* strain in stationary phase, with our *in vitro* data where HelD removes RNAP from promoter DNA, and with our model, in which HelD stochastically displaces RNAPs from stationary phase promoters (Fig. 7C). This is also consistent with the dampening effect of HelD on *in vitro* transcription reported previously (Hurst-Hess *et al*., 2022; Kovaľ *et al*., 2024).

Since G is the predominant initiation nucleotide in *M. smegmatis* (Fig. 5A), the smaller pool of GTP in stationary phase might also contribute to the eventual prolonged periods of the promoter-associated RNAPs. Moreover, both exponential and stationary phase genes are characterized by G being present at +1 position in approximately 50 % of genes (Fig. 5A). This possibly implies that the decreased GTP level in stationary phase might affect the whole genome transcription initiation kinetics. As stationary phase transcription initiation complexes wait for GTP, the RNAP molecules within these complexes might become targets of HelD-mediated release. Another important point is that HelD has been demonstrated to hydrolyze GTP *in vitro* and that this step is required for the release of HelD-bound RNAP (Kouba *et al*., 2020). In stationary phase, if HelD is present, lower GTP level might contribute to a decreased efficiency of RNAP release, making it unavailable for the next rounds of transcription.

Based on recently published cryo-EM structures, HelD binds RNAP initiation complexes without any specific additional protein factor (Kouba *et al*., 2020; Kovaľ *et al*., 2024). Therefore, we assume that HelD stochastically removes RNAPs from promoters. Intriguingly, HelD binding to RNAP is incompatible with RNAP’s association with CarD (Kovaľ *et al*., 2024), and we do not detect CarD and HelD in the same RNAP complex (Fig. 7A). Based on cryo-EM, CarD and HelD binding to RNAP is mutually exclusive (Kovaľ *et al*., 2024). We cannot exclude that CarD protects the RNAP initiation complex at the promoter from being removed by HelD (Fig. 7B). In addition, both CarD and HelD are detected by ChIP-seq on the same promoters. As CarD is predominantly present in exponential phase cells as a free protein, not in complex with RNAP core or holoenzyme (Fig. 6G), we propose that the association of CarD with the transcription initiation complex might be transient and/or short-lived. Nevertheless, our ChIP-seq experiments lead us to the conclusion that a subset of *M. smegmatis* promoters can accommodate HelD, RbpA and CarD as well, although probably not necessarily all three transcription factors within the same complex. HelD and CarD can bind to the same promoters, but their binding is probably sequential, and they cannot be found at the same promoter at the same time.

As determined previously, HelD was significantly upregulated at both RNA and protein levels when *M. smegmatis* cells were treated with sublethal concentrations of rifampicin (Giddey *et al*., 2017; Hurst-Hess *et al*., 2019) and it was demonstrated to convey the rifampicin resistance in *M. smegmatis, M. abscesus* and *S. venezuelae* (Hurst-Hess *et al*., 2022; Surette *et al*., 2022). We showed for the first time that HelD contributes to rifampicin resistance *in vivo* in liquid cultures of *M. smegmatis.* HelD presence also partially correlates with improved growth in rifampicin in many clinically important mycobacterial species. Mycobacteria significantly differ in their sensitivity to rifampicin; for example, *Mycobacterium avium* is able to grow at the highest rifampicin concentration tested. In contrast, less than 5 % of *M. tuberculosis* clinical isolates were resistant to rifampicin and this resistance was mainly caused by amino acid substitutions in the RNAP β-subunit, which interfere with rifampicin binding (Dohál *et al*, 2022). Nevertheless, RNAP β-subunit mutations that confer rifampicin resistance reduce the fitness of *M. tuberculosis* and, therefore, are not widespread (Gygli *et al*, 2017). The differences in rifampicin resistance among species are rather caused by the presence of specific genes. The presence of the *helD* gene appears to be a reliable predictor of growth advantage in the presence of rifampicin, but our findings also strongly suggest the additional resistance mechanisms among mycobacterial species. *M. avium* is resistant to 50 μg/ml rifampicin despite lacking both *helD* and *arr* genes. This highlights the potential existence of other, yet unidentified, genes associated with rifampicin resistance in mycobacteria, which could, for example, involve efflux pumps that contribute to antibiotic insensitivity (Adams *et al*, 2011).

We have shown that even in the absence of rifampicin, *M. smegmatis* cells benefit from HelD, especially when it comes to exponential growth associated transcription of highly expressed genes like those encoding tRNAs. We propose that HelD functions as a transcriptional regulatory protein in rapidly growing mycobacteria, with its role in rifampicin resistance potentially being a secondary effect of its primary function. Proteins with such dual functions should be considered in future studies of bacterial antibiotic resistance mechanisms. Future studies will reveal compounds to block HelD function leading to a possible generation of therapeutic agents useful against non-tuberculous mycobacteria.

## MATERIALS AND METHODS

For the information on oligonucleotides used in this manuscript, please see Supplementary Material (Supplementary Table S2).

### Cloning of plasmids, bacterial strains

The *MSMEG_2174* (*helD*) deletion strain was generated in the following way. First, the cassette for single crossing over was generated by combining pUC18 plasmid digested with BamHI and HindIII with three PCR fragments amplified by Q5 High-Fidelity DNA Polymerase (NEB): 500 bp of the left homology arm (sequence located upstream of the *helD* coding sequence, *M. smegmatis* mc^2^ 155 chromosomal DNA as a PCR template), a hygromycin resistance encoding sequence (Huff *et al*, 2010; Šiková *et al*., 2019) and a 500 bp of the right homology arm (sequence located downstream of the *helD* coding sequence, *M. smegmatis* mc^2^ 155 chromosomal DNA as a PCR template). The four DNA fragments were assembled using a Gibson assembly cloning kit (NEB). The mixture was transformed into DH5α competent *E. coli* cells (NEB) and the resulting construct was verified by sequencing. The fragment containing the cassette for homologous recombination was subsequently transformed into the *M. smegmatis* pJV53 strain [increased frequency of homologous recombination, (van Kessel & Hatfull, 2007)] and individual clones were selected for hygromycin and kanamycin resistance. Subsequently, the clones were cured of pJV53, and one clone was selected, which was hygromycin resistant and kanamycin sensitive (LK2650) and was used in further experiments.

Each of the *M. smegmatis* strains used for the expression of C-terminally FLAG-tagged CarD (*MSMEG_6077*), RbpA (*MSMEG_3858*) and HelD (*MSMEG_2174*) carried an extra copy of the respective protein. The coding sequences of *MSMEG_6077*, *MSMEG_3858* and *MSMEG_2174* were amplified by PCR with *M. smegmatis* mc^2^ 155 chromosomal DNA as a template and the FLAG tag sequence was encoded in the reverse primers. Subsequently, these PCR fragments were inserted into the pTet-Int plasmid (Choudhary *et al*, 2015) via NdeI and HindIII, transformed into *E. coli* DH5α, sequence verified, and the constructs were integrated into *M. smegmatis* mc^2^ 155 genome. The expression of FLAG-tagged CarD, RbpA and HelD was controlled by the addition of anhydrotetracycline (ATc) to the growth media at a concentration of 10 ng/ml (see section Bacterial growth, rifampicin treatment). The inducible RbpA-FLAG strain and the inducible HelD-FLAG strain have been described previously (Kouba *et al*., 2020).

The strain with FLAG tag knock-in in the native *MSMEG_1368* (*rpoC*) locus was generated in the following way. Final construct consisted of a hygromycin resistance cassette (HYG) flanked by left and right arms [left arm (LA) - 500 bp long region homologous to the 3’ terminal part of the gene and containing the sequence encoding the affinity tag (1x FLAG-tag: DYKDDDDK); right arm (RA) - 500 bp long region homologous to the 3’ end of the targeted gene]. DNA fragments (LA, HYG, RA) were amplified by PCR with Q5 High-Fidelity DNA Polymerase (NEB). *M. smegmatis* mc^2^ 155 chromosomal DNA and plasmid containing HYG (Huff *et al*., 2010; Šiková *et al*., 2019) served as templates. Fragments were assembled into the pUC18 plasmid cloning vector using Gibson Assembly Cloning Kit (NEB). The mixture was transformed into DH5α competent *E. coli* cells (NEB) and the resulting construct was verified by sequencing, then cleaved with restriction enzymes BamHI and HindIII and the linear cassette for homologous recombination was transformed by electroporation into *M. smegmatis* electrocompetent cells containing the pYS1 integrating plasmid vector (Shenkerman *et al*, 2014). Individual clones were selected for hygromycin and kanamycin resistance. Then, the clones were cured of pYS1 plasmid, resulting in final *M. smegmatis* strain LK2740 (RNAP- FLAG) with 1x FLAG-tag incorporated at the respective native locus. This strain was utilized in RNAP protein level quantification (western blot).

The HelD purification strain has been described previously (Kouba *et al*., 2020). The CarD and RbpA expression vectors were generated by PCR cloning using vector pET302/NT-His, verified by sequencing and transformed to *E. coli* DE3 cells to generate CarD and RbpA purification strains. These three strains were utilized in CarD, RbpA and HelD protein purification, the protein level quantification (western blot), and antibody production.

### Bacterial growth, rifampicin treatment

*Mycobacterium smegmatis* mc^2^ 155 cells were grown at 37 °C in Middlebrook 7H9 medium with 0.2% glycerol and 0.05% Tween 80 and harvested in exponential (OD_600_ ∼0.5) or early stationary phase (OD_600_ ∼2.5–3, 24 hours of cultivation). Rifampicin was added at OD_600_ ∼0.1, and cells were grown for 6-7 hours. The CarD, RbpA and HelD-FLAG tagged strains were incubated with 10 ng/ml ATc for 3 hours (ATc was added after 3 hours of growth, during exponential phase) before being harvested for the downstream procedures (western blot, ChIP-seq). For the induction in stationary phase, ATc was added 14 hours before harvesting the cells for the downstream procedures (western blot). For determination of growth curve in Fig. 2A, 300 μl of 7H9-ADC medium containing 0.5% (vol/vol) glycerol, 0.05% tyloxapol with and without rifampicin was pipetted into individual wells of a 100-well polystyrene honeycomb plate. 10 μl of the exponentially grown *M. smegmatis* culture was used for inoculation to final OD_600_ 0.05. Plates were cultivated at 37 °C and 400 rpm and OD value recorded at 15 min intervals. Three independent biological replicates were performed. The MIC values for wt and Δ*helD* strains were determined by resazurin assay described in (Knejzlík *et al*, 2020). Briefly, rifampicin was serially two-fold diluted in the range of 50 – 0 μg/ml in 100 μl 7H9-ADC medium containing 0.5% (vol/vol) glycerol and 0.05% tyloxapol in 96-well plate. Individual wells were inoculated by 5 μl culture in the exponential phase of growth to final OD_600_ 0.005. Plates were incubated at 37°C for 24 hours, 15 μl of a 0.02% resazurin solution in phosphate-buffered saline (PBS) was added, and the plate was incubated for an additional 24 hours.

Patient-derived isolates were cultivated following the Clinical and Laboratory Standards Institute (CLSI) guidelines (Woods *et al*, 2011) on Löwenstein–Jensen using the following rifampicin concentrations: 20 μg/ml and 50 μg/ml. Incubation was carried out at 37 °C for 3 to 7 days for rapidly growing mycobacteria and 3 to 6 weeks for slow-growing mycobacteria.

### SDS-PAGE, western blotting

Cells were harvested by centrifugation, pellets washed with lysis buffer [20 mM Tris–HCl pH 7.9, 150 mM KCl, 1 mM MgCl_2,_ 1 mM dithiothreitol (DTT), 0.5 mM phenylmethylsulfonyl fluoride (PMSF) and Calbiochem Protease Inhibitor Cocktail Set III protease inhibitors] and lysed by sonication (Sonopuls HD3100, Bandelin [Germany]; 50 % amplitude 15 × 10 s with 1 min pauses on ice), and centrifuged at 8960 x g for 15 min at 4 °C. The proteins of interest were resolved by SDS-PAGE (Nu-PAGE, 4-12% Bis-Tris precast gels, Invitrogen) and stained with Coomassie. Proteins were detected by western blotting using an anti-RNAP β subunit antibody [clone 8RB13] (BioLegend), anti-σ^70^ antibody [clone 2G10] (BioLegend), anti-HelD antibody, anti-CarD antibody, anti-RpbA antibody and an HRP-labeled anti-mouse IgG antibody (Sigma-Aldrich). The anti-HelD, CarD and RbpA antibodies are in-house produced. For detailed information, see the section Antibodies production. The purified proteins used for standard curves in protein level quantification were prepared as described in the following section,

Protein purification. We calculated the amounts of RNAP, CarD, RbpA and HelD, expressed in nanograms (ng) per microgram (µg) of lysate and further normalized these values to the molecular weight of each protein to determine RNAP, CarD, RbpA and HelD molecular ratios in the cell lysate.

Purified σ^A^ and RNA polymerase used for protein level quantification (western blot) were kindly provided by Barbora Brezovská and Nabajyoti Borah, respectively.

### Immunoprecipitation

For immunoprecipitation, cells were pelleted and washed in lysis buffer (20 mM Tris–HCl pH 7.9, 150 mM KCl, 1 mM MgCl_2_), pelleted again and these pellets were frozen at -70 °C. Bacterial pellets were resuspended in lysis buffer supplemented with PMSF and Protease inhibitor cocktail [20 mM Tris–HCl pH 7.9, 150 mM KCl, 1 mM MgCl_2_, 1 mM dithiothreitol (DTT), 0.5 mM PMSF, Protease Inhibitor Cocktail Set III protease inhibitors (Calbiochem)], sonicated 15 × 10 s with 1 min pauses on ice and centrifuged at 8960 x g for 15 min at 4 °C. 1 mg of protein lysates were incubated for 16-18 h at 4 °C with 25 μl of M2 anti-FLAG resin (Sigma Aldrich). The captured complexes were washed four times using 20 mM Tris–HCl pH 7.9, 150 mM KCl, 1 mM MgCl_2_. The immunoprecipitated proteins were eluted using 60 µl of Tris buffer saline (50mM Tris-HCl pH 7.5, 150 mM NaCl) containing 150 ng/ml of FLAG peptide for 30 minutes at 4 °C, analyzed on SDS-PAGE (Nu-PAGE, 4-12% Bis-Tris precast gels, Invitrogen) and stained with Coomassie.

### Protein purification and antibodies production

*E. coli* DE3 strains were grown at 37 °C in LB medium and shaken at 120 rpm for ∼ 3 h. Protein expression was induced at OD_600_ ∼0.6 with 0.8 mM IPTG (for CarD and RbpA) or 0.5 mM IPTG (for HelD) for 3 hours at room temperature. The cultures were cooled and centrifuged at 6000 x g for 10 min at 4 °C. The pellets were washed with T-buffer (300 mM NaCl, 50 mM Tris-HCl pH 7.5, 5% glycerol) and stored at -70 °C. Bacterial pellets were resuspended in T-buffer (supplemented with 3 mM β-mercaptoethanol), sonicated (Sonopuls HD3100, Bandelin [Germany]; 50 % amplitude, 15 x 10 s pulse, 1 min pause on ice) and centrifuged at 27000 x g, 4 °C (Beckman Coulter, JA 25-50 rotor) for 10 min. The lysates were incubated with 1 ml of prewashed Ni-NTA Agarose beads (QIAGEN) for 1.5 h at 4 °C. Ni-NTA beads with bound proteins were centrifuged at 2000 rpm, 4 °C for 5 min, washed with 10 ml T-buffer, then washed with 10 ml T-buffer containing 30 mM imidazole, resuspended in 500 μl of T-buffer with 400 mM imidazole and serial fractions (500 μl) of the eluted proteins were collected. Their concentrations and purity were determined by Bradford assay and on SDS-PAGE, respectively. The selected fractions containing HelD/CarD/RbpA proteins were pooled and dialyzed for 20 h against TEV Cleavage Buffer (50 mM Tris-HCl pH 8, 100 mM NaCl, 0.5 mM EDTA, 0.5 mM DTT, 5% glycerol) in Slide-A-Lyzer Dialysis Cassette 3.500 MWCO (Thermo Scientific). The TEV protease was added to the dialyzed proteins at a TEV protease:protein ratio of 1:20. The cleavage was performed for 8-12 hours at 4 °C.

The cleaved protein samples were incubated with 1 ml of prewashed Ni-NTA Agarose beads for 1.5 h at 4 °C to remove the cleaved NT-His tags and His-tagged TEV protease from protein solutions. Ni-NTA-protein suspensions were then centrifuged at 3500 x g, 4 °C for 3 min and the supernatants containing the cleaved proteins were transferred into the new tube. The Ni-NTA beads were further washed (2x) with 1 ml TEV Cleavage Buffer and the supernatants were transferred into new tubes. The Ni-NTA-bound His-tags were eluted with T-buffer (400 mM imidazole) to check the protein-His tag cleavage.

Cleaved protein samples (supernatants) and the eluate were visualized and checked by SDS-PAGE stained with Coomassie. Individual cleaved protein fractions (of HelD or CarD or RbpA) were mixed and dialyzed against storage buffer (50 mM Tris-HCl, pH 8, 100 mM NaCl, 50% glycerol, 3 mM β-mercaptoethanol) and stored at -20 °C. Protein concentrations were measured by Qubit.

### Animal experiments and immunizations

All animal experiments were approved by the Animal Welfare Committee of the Institute of Molecular Genetics of the Czech Academy of Sciences, v. v. i., in Prague, Czech Republic. Handling of animals was performed according to the Guidelines for the Care and Use of Laboratory Animals, the Act of the Czech National Assembly, Collection of Laws no. 246/1992. Permission no. 19/2020 was issued by the Animal Welfare Committee of the Institute of Molecular Genetics of the Czech Academy of Sciences in Prague.

Five-week-old female BALB/cByJ mice (Charles River, France) were immunized by intraperitoneal injection with proteins HelD, RbpA or CarD (20 μg in 200 μl) adjuvanted with aluminum hydroxide (Alum, SevaPharma, Czech Republic). Mice received three doses of the protein in 2 weeks intervals and 1 week after the third immunization, the blood was collected from anesthetized animals (i.p. injection of 80 mg/kg ketamine and 8 mg/kg xylazine) by retroorbital puncture method. Sera were recovered from the supernatant after centrifugation of clogged blood at 5000 x g for 10 min at 8 °C and stored at –20 °C. The specificity of the antibodies was checked with 1 μg of *M. smegmatis* cell lysate with 1:5000 antibody dilution.

### Glycerol gradient ultracentrifugation

*M. smegmatis* exponential and stationary phase cells were pelleted and resuspended in 20 mM Tris–HCl pH 8, 150 mM KCl, 1 mM MgCl_2_, 1 mM dithiothreitol (DTT), 0.5 mM phenylmethylsulfonyl fluoride (PMSF) and Calbiochem Protease Inhibitor Cocktail Set III protease inhibitors, sonicated 15 × 10 s with 1 min pauses on ice and centrifuged. Protein extracts (1 mg) were loaded on a linear 10–30% glycerol gradient prepared in gradient buffer (20 mM Tris–HCl pH 8, 150 mM KCl, 1 mM MgCl_2_) and fractionated by centrifugation at 32 000 rpm (130 000 × g) for 17 h using an SW-41 rotor (Beckman). The gradient was divided into 20 fractions and proteins from each fraction were resolved on SDS-PAGE and detected by western blotting.

### ChIP-seq

For the RNAP and σ^A^/σ^B^ exponential and stationary phase ChIP-seq, the experiment was performed as described previously (Vanková Hausnerová *et al*., 2024), using *M. smegmatis* wild type strain. For the CarD, RbpA and HelD exponential phase ChIP-seq, the published protocol was modified in the following way. 100 ml of bacterial culture was crosslinked with 1% formaldehyde (final concentration) for 30 min at 37 °C in the shaker. Formaldehyde was quenched by glycine (0.125 M final concentration) added for 5 min at 37 °C in the shaker. Bacterial cultures were centrifuged for 5 min at 8960 x g, 4 °C, the pellet was washed with 10 ml of 1x PBS, bacteria were centrifuged again and the pellet was immediately frozen at -70 °C. The bacterial pellet was resuspended in 3 ml of ice-cold RIPA buffer (150 mM NaCl, 1% Triton X-100, 0.5% deoxycholate, 0.1% SDS, 50 mM Tris-HCl pH 8.0, 5 mM EDTA) with protease inhibitors cocktail (5 μl/ml, Sigma-Aldrich) and 0.5 mM PMSF and sonicated 18 x 10 seconds with 1 min pauses on ice between cycles (Sonopuls HD3100, Bandelin [Germany]; 50 % amplitude). The lysate was centrifuged at 8960 x g and 4 °C for 15 min, the pellet was discarded. A volume corresponding to 2 mg of protein was incubated with 20 μl of M2 anti-FLAG resin (Sigma Aldrich). The captured complexes were washed twice with RIPA buffer (150 mM NaCl, 1% Triton X-100, 0.5% deoxycholate, 0.1% SDS, 50 mM Tris-HCl pH 8.0, 0.5 mM EDTA), four times with LiCl buffer (100 mM Tris-HCl pH 8.5, 500 mM LiCl, 1% Triton X-100, 1% deoxycholate), two times with RIPA and twice with TE buffer (10 mM Tris-HCl pH 8.0, 1 mM EDTA). Protein-DNA complexes were eluted with elution buffer (50 mM Tris-HCl pH 8, 0.66 mM EDTA, 1 % SDS) for 10 min at 65 °C, decrosslinked in the presence of 200 mM NaCl for 5 h at 65 °C, treated with 100 µg/ml RNase A for 1 h at 37 °C and 400 µg/ml proteinase K for 30 min at 45 °C. DNA was purified with the QIAGEN PCR purification kit and eluted with 100 μl of Elution Buffer. 40 µl of immunoprecipitated DNA sample or 10 ng of DNA input were used for library construction according to the NEXTflex™ ChIP-Seq Kit manual including the Size-Selection Cleanup step B1. The libraries were checked using Agilent 2100 Bioanalyzer High Sensitivity DNA Assay Chip. Pooled barcoded libraries (one biological triplicate) were sequenced in single lanes using the Illumina NextSeq® 500/550 High Output Kit v2 in 75 bp single end regime at the Institute of Molecular Genetics AS CR, Prague, Czech Republic.

For ChIP results quantification, the captured DNA was amplified by RT-qPCR in a LightCycler 480 System (Roche Applied Science) in duplicate reactions containing LightCycler 480 SYBR Green I Master and 0.5 μM primers (each). Primers were designed with Primer3, sequences are listed in the Supplementary Data. Negative controls (no template reactions) were run in each experiment, the quality of the PCR products was determined by dissociation curve analysis and the efficiency of the primers was determined by standard curves. The relative amounts of co-immunoprecipitated DNA were quantified on the basis of threshold cycles (Ct) for each PCR product that was normalized to input values according to the formula 2(Ct^(immunoprec)^ − Ct^(input)^).

### RNA-seq

Before total RNA extraction, an mRNA spike-in mix composed of four different eukaryotic mRNAs was added [Plat, Moc, Elav2 and nLuc; the mRNAs were prepared by *in vitro* transcription from pJET plasmids using the MEGAscript T7 Transcription Kit (Thermo Fisher Scientific), the sequences of the mRNAs are provided in the Supplementary Material]. The amount of RNA spike-ins was 10 ng per 30 ml of culture at an OD_600_ of 0.5. Each frozen cell pellet was resuspended in 240 μl TE buffer (pH 8.0) plus 60 μl LETS buffer (50 mM Tris-HCl pH 8.0, 500 mM LiCl, 50 mM EDTA pH 8.0, 5% SDS) and 600 μl acidic (pH∼3) phenol/chloroform (1:1). Lysates were sonicated in a fume hood, centrifuged, the aqueous phase extracted two more times with acidic phenol/chloroform and precipitated with ethanol. RNA was dissolved in double distilled water and treated with DNase (TURBO DNA-free Kit, Ambion). 1 μg of DNase-treated RNA was ribodepleted (riboPOOL Kit Pan-Actinobacteria, siTOOLs). The sample integrity was checked using Agilent 2100 Bioanalyzer Pico Chip. The ribodepleted RNA sample (20-100 ng) was used for library construction according to the NEXTFLEX® Rapid Directional RNA-Seq Kit. The libraries were checked using Agilent 2100 Bioanalyzer High Sensitivity DNA Assay Chip. Pooled barcoded libraries were sequenced in a single lane with Illumina NextSeq® 500/550 High Output Kit v2 in 75 bp single end regime at the Institute of Molecular Genetics AS CR, Prague, Czech Republic.

For RNA-seq validation, 5 μl RNA (∼ 2.5 μg) was reverse transcribed into cDNA (20 μl reaction, SuperScriptIII, Invitrogen) using random hexamers and amplified by RT-qPCR in a LightCycler 480 System (Roche Applied Science). The mRNA level was normalized to the value of the Plat mRNA spike-in according to the formula 2^(Ct^(spike)^ − Ct^(mRNA)^) and expression (E) normalized to the nontreated strain (E =E_deletion strain_/E_wildtype strain_, alternatively E =E_treated sample_/E_non-treated sample_).

### NGS data analysis

#### RNA-seq

Read quality was checked using FastQC version 0.11.9 (https://www.bioinformatics.babraham.ac.uk/projects/fastqc/). When needed, adapters and low-quality sequences were removed using Trimmomatic 0.39 (Bolger *et al*, 2014). Reads were aligned to the reference genome using HISAT2 2.2.1 (Kim *et al*, 2015) and SAMtools 1.13 (Bonfield *et al*, 2021; Li *et al*, 2009). Read coverage tracks were computed using deepTools 3.5.1 (Ramírez *et al*, 2016). The DESeq2 R package (Love *et al*, 2014) was used to identify differentially expressed genes at FDR ≤ 0.05. For the detection of transcriptome-wide shifts in cellular RNA content, a global normalization using a mix of four spike-in control RNAs (Plat, Mos, Elav2, Nluc) was performed.

#### ChIP-seq

ChIP-seq peak calling and gene assignment was done as described in (Vanková Hausnerová *et al*., 2024). Briefly, the reads were mapped to *M. smegmatis* genome (NCBI RefSeq NC_008596.1) with HISAT2 (Kim *et al*., 2015) and peaks were called by MACS2 (Zhang *et al*, 2008) on each replicate separately and only the peak regions overlapping in all replicates were retained as resulting final peaks. The highest (worst) p-value from the overlapping peaks was assigned to the resulting peak as its p-value. The lowest -log_10_q-value from three replicas is shown in Supplementary Table S1 (the higher the -log_10_q-value, the higher the statistical significance of the peak). Peaks with -log_10_q-value>40 were used for further analysis. The BEDTools suite v2.30.0 was used for peak annotation and analysis (Quinlan & Hall, 2010). The Venn diagrams for overlapping peaks were created with BEDTools intersect (Quinlan & Hall, 2010) and matplotlib-venn package. The 3-way Venn diagram was created with Intervene (Khan & Mathelier, 2017). The coverage profiles of the 1 kb region around ORF start were created with deepTools (computeMatrix, plotProfile) (Ramírez *et al*., 2016) and programming libraries rtracklayer (Lawrence *et al*, 2009), pandas (McKinney, 2010), matplotlib (Hunter, 2007), seaborn (Waskom *et al*, 2020).

#### HelD and Arr homologs in 15 mycobacterial species

To check whether a species contains homologous sequence for HelD (gene *MSMEG_2174*) or Arr (gene *MSMEG_1221*) protein, we have obtained proteins defined in genomes from GenBank (Benson *et al*, 2013) with the same taxonomy id using the NCBI-datasets command line interface (O’Leary *et al*, 2024). Then we used BLAST (Altschul *et al*, 1997) to find homologous proteins; we considered only those hits with E-value less than 10^-25^ and bitscore more than 145 for Arr and 734 for HelD (½ of max score respectively). If homolog was found in more than 80 % of the genomes of the species, we considered the species to have homologous protein. Association between prevalence of resistance and presence of *helD*/*arr* homolog in published genomes was evaluated with a binomial generalized linear mixed model using the lme4 package (Bates *et al*, 2015) - the presence of the homolog in the strain was treated as a fixed effect and the model included a varying intercept for each strain. P-values were derived from a likelihood ratio test between the full model and a model with just the strain varying intercept.

#### Initiating nucleotides frequency

To assess frequency of nucleotides at and downstream of TSS, we have used primary TSS (pTSS) (Martini *et al*, 2019) for genes with detectable (non-zero) expression in exponential and stationary phase in the wt *M. smegmatis* (Šiková *et al*., 2019). For each gene with defined pTSS and detectable expression we have extracted first 10 nucleotides (TSS included) and used WebLogo (Crooks *et al*, 2004) to plot distribution of nucleotides at each position.

#### Nucleotide metabolomic analysis

Stationary phase of growth was achieved as follows: the 50 ml 7H9 medium containing 0.2% glycerol and 0.05% Tyloxapol was inoculated with a fresh culture of *M. smegmatis* to an initial OD_600_ ∼0.1 and cultures were grown at 37 °C and 200 rpm. Exponential and stationary phase was achieved at OD_600_ ∼0.4 (4.5 h) and ∼5 (18 h), respectively. 5 ml (exponential phase) or 1 ml (stationary phase) of the cell suspension was quickly vacuum filtered through a 0.45 μm/25 mm cellulose acetate filter. The membrane with the collected bacteria was immediately transferred into 1 ml of ice-cold 1 M acetic acid in a 1.5 ml Eppendorf tube and quickly frozen in liquid nitrogen. Nucleotide content was determined as is described previously (Knejzlík *et al*, 2022) and expressed in nmol per OD_600_ unit.

### *In vitro* RNAP release assay

First, the open complex promoter bubble was formed by annealing 15 μl of each oligonucleotide (100 μM stock concentration) in a thermocycler with the following program: 98 °C for 5 min, 95 °C for 1 min, then the temperature decreased by 1 °C every 1 min for 70 cycles. The non-template strand + template strand oligonucleotides (Kovaľ *et al*., 2024) were annealed, one pair of oligos with the biotinylated 5′end of the non-template strand oligo and the other pair of oligos without any biotin, which served as competitor DNA. The open promoter complexes were reconstituted in 10 μl of transcription buffer (40 mM Tris-HCl, pH 8.0, 10 mM MgCl_2_, 1 mM DTT), one reconstitution contained 8.8 pmol RNAP, 12.5 pmol σ^A^ and 5 pmol of the annealed pair of oligonucleotides with the biotinylated 5′end of the non-template strand. After the incubation for 5 min at 25 °C with 260 rpm shaking, the complexes were mixed with 5 μl of streptavidin coated magnetic beads (Dynabeads MyOne Streptavidin C1, ThermoFischer; the beads were washed with transcription buffer before mixing with the reconstitutions). The open promoter complexes were bound to the beads for 10 min at 25 °C with 260 rpm shaking. Subsequently, the beads were separated using a magnetic rack. The supernatant was removed. The beads with bound open promoter complexes were washed and incubated with 10 μl of transcription buffer containing 40 pmol of competitor DNA and 15 pmol of HelD for 10 min at 25 °C with 260 rpm shaking. HelD was not included in the control reactions. Next, the beads were again separated using the magnetic rack, the supernatant containing the released proteins was saved and the beads were washed with transcription buffer. Both the supernatants and the beads were resolved on SDS-PAGE gels and visualized by staining with Coomassie.

The RNAP protein used in this experiment was purified as described previously (Kouba *et al*, 2019). The σ^A^ protein was kindly provided by Barbora Brezovská. The HelD protein was purified according to the protocol described in the section Protein purification and antibodies production (see above).

## DATA AVAILABILITY

- Data and code for the analysis: Zenodo (10.5281/zenodo.11110137)
- RNA-seq analysis of *M. smegmatis* wild type and *helD* deletion strains: BioStudies ArrayExpress E-MTAB-13810 (https://www.ebi.ac.uk/biostudies/arrayexpress/studies/E-MTAB-13810)
- *M. smegmatis* HelD ChIP-seq: BioStudies ArrayExpress E-MTAB-11827 (https://www.ebi.ac.uk/biostudies/arrayexpress/studies/E-MTAB-11827)
- *M. smegmatis* CarD ChIP-seq: BioStudies ArrayExpress E-MTAB-11828 (https://www.ebi.ac.uk/biostudies/arrayexpress/studies/E-MTAB-11828)
- *M. smegmatis* RbpA ChIP-seq: BioStudies ArrayExpress E-MTAB-11829 (https://www.ebi.ac.uk/biostudies/arrayexpress/studies/E-MTAB-11829)

## ACKNOWLEDGEMENTS

We would like to thank Barbora Brezovská and Nabajyoti Borah for the kind donation of purified proteins and Tomáš Kouba for critical reading of the manuscript.

This work was supported by the Czech Science Foundation grant No. 23-05622S (to J. Hn.), The Charles University Grant Agency grant No. 275823 to M. Sh., National Institute of virology and bacteriology (Program EXCELES, ID Project No. LX22NPO5103) - Funded by the European Union - Next Generation EU (to Z. K. and H. Š.), OPJAK Talking microbes - understanding microbial interactions within One Health framework (CZ.02.01.01/00/22_008/0004597 to O.S.), ELIXIR CZ research infrastructure project (MEYS Grant No. LM2023055) including access to computing and storage facilities (to M. Sch. and M. M.), Ministry of Health - conceptual development of research organization [The National Institute of Public Health (NIPH), 75010330] and by the Czech National Node to the European Infrastructure for Translational Medicine (project No. LM2023053) from the Ministry of Education, Youth and Sports of the Czech Republic (to O. S.).

## AUTHOR CONTRIBUTIONS

Conceptualization, J. Hn., D. K., Z. K., V. D. and V. V. H.; Methodology, D. K., M.P., M. Sch., Z. K., V. D., V. V. H., and J. Hn.; Software, M. P., M. M. and M. Sch.; Investigation, D. K., V. V. H., M. Sh., S. N., J. J. M., J. Ha., M. Š., D. Ká., P. H., H. Š., J. Ho., M. D., O. S., Z. K., V. D., and J. Hn.; Data Curation, M. P.; Writing – Original Draft V. V. H. and J. Hn.; Visualization: V. V. H., M. Sh., M. M. and J. Hn., Supervision: J. Hn.; Funding Acquisition: J. Hn., Z. K., L. K. and O. S.

## CONFLICT OF INTEREST STATEMENT

The authors declare no competing interests.

